# Myosin 18Aα targets guanine nucleotide exchange factor β-Pix to dendritic spines of cerebellar Purkinje neurons to promote spine maturation

**DOI:** 10.1101/2020.02.12.944488

**Authors:** Christopher J. Alexander, Melanie Barzik, Ikuko Fujiwara, Kirsten Remmert, Ya-Xian Wang, Ronald S. Petralia, Thomas B. Friedman, John A. Hammer

## Abstract

Dendritic spines are signaling microcompartments that serve as the primary site of synapse formation in neurons. Actin assembly and myosin 2 contractility play major roles in the maturation of spines from filopodial precursors, as well as in the subsequent, activity-dependent changes in spine morphology that underly learning and memory. Myosin 18A is a myosin 2-like protein conserved from flies to man that lacks motor activity, is sub-stochiometric to myosin 2, and co-assembles with myosin 2 to make mixed filaments. Myosin 18A is alternatively spliced to create multiple isoforms that contain unique N- and C-terminal extensions harboring both recognizable and uncharacterized protein: protein interaction domains. These observations suggest that myosin 18A serves to recruit proteins to mixed filaments of myosin 2 and myosin 18A. One protein known to bind to myosin 18A is β-Pix, a guanine nucleotide exchange factor (GEF) for Rac1 and Cdc42. Notably, β-Pix has been shown to promote spine maturation by activating both Arp2/3 complex-dependent branched actin filament assembly and myosin 2 contractility within spines. Here we show that myosin 18Aα is expressed in cerebellar Purkinje neurons and concentrates in spines along with myosin 2 and F-actin. Myosin 18Aα is targeted to spines by co-assembling with myosin 2 and by an actin binding site present in its N-terminal extension. miRNA-mediated knockdown of myosin 18Aα results in a significant defect in spine maturation that is rescued by an RNAi-immune version of myosin 18Aα. Importantly, β-Pix co-localizes with myosin 18Aα in spines, and its spine localization is lost upon myosin 18Aα knockdown or when its myosin 18Aα binding site is deleted. Finally, we show that the spines of myosin 18Aα knockdown Purkinje neurons contain significantly less F-actin and myosin 2. Together, these data demonstrate that mixed filaments of myosin 2 and myosin 18Aα form a complex with β-Pix in Purkinje neuron spines that promotes spine maturation by enhancing the assembly of actin and myosin filaments downstream of β-Pix’s GEF activity.

## INTRODUCTION

Dendritic spines are small protrusions on the surface of neuronal dendrites that form the postsynaptic component of most excitatory synapses in the brain, function as signaling microcompartments, and serve as the primary site of memory formation (Yuste and Bonhoeffer, 2001; Okamoto *et al*., 2004; Lamprecht and LeDoux, 2004; Matsuzaki *et al*., 2004; Bourne and Harris, 2008; Holtmaat and Svoboda, 2009; Newpher and Ehlers, 2009; Roberts *et al*., 2010; Kasai *et al*., 2010; Fortin, Srivastava and Soderling, 2012; Hlushchenko, Koskinen and Hotulainen, 2016; Basu and Lamprecht, 2018; Nakahata and Yasuda, 2018). Actin filaments comprise the major structural element of spines, and changes in spine size and shape that occur upon synapse usage are thought to result largely from changes in spine actin content, organization and dynamics (Yuste and Bonhoeffer, 2001; Lamprecht and LeDoux, 2004; Matsuzaki *et al*., 2004; Okamoto *et al*., 2004; Bourne and Harris, 2008; Holtmaat and Svoboda, 2009; Newpher and Ehlers, 2009). Moreover, these actin-dependent changes in spine morphology (structural plasticity) are thought to promote the long-lasting changes in synaptic strength (functional plasticity) that underlie learning and memory (Kasai *et al*., 2010; Roberts *et al*., 2010; Fortin, Srivastava and Soderling, 2012; Hlushchenko, Koskinen and Hotulainen, 2016; Basu and Lamprecht, 2018; Nakahata and Yasuda, 2018). Consistently, many learning and memory disorders (e.g. autism, psychosis) are associated with actin-related defects in dendritic spine morphology and dynamics (Yan *et al*., 2016; Lima Caldeira, Peça and Carvalho, 2019). Defining the molecular basis of learning and memory in both health and disease will require, therefore, a thorough understanding of the origin, organization and dynamics of actin in spines.

While several models have been proposed for how dendritic spines are created, the prevailing model is that they begin life as dendritic filopodia, and that these filopodia then mature into spines (Yuste and Bonhoeffer, 2004; Ethell and Pasquale, 2005; Tada and Sheng, 2006; Yoshihara, De Roo and Muller, 2009; Kanjhan, Noakes and Bellingham, 2016). Like conventional filopodia, dendritic filipodia contain long, unbranched actin filaments (Korobova and Svitkina, 2010) created by a formin (in this case, mDia2, downstream of the Rho-related GTPase Rif) (Hotulainen *et al*., 2009). Unlike conventional filopodia, dendritic filopodia also contain branched actin filaments (Korobova and Svitkina, 2010). Whether these branched filaments are created by the Arp2/3 complex is not entirely clear (Korobova and Svitkina, 2010; Spence *et al*., 2016). What is much clearer, however, is that the maturation of dendritic filopodia into dendritic spines involves a large increase in the creation of branched actin filaments by the Arp2/3 complex (Kim *et al*., 2006; Soderling *et al*., 2007; Haeckel *et al*., 2008; Wegner *et al*., 2008; Spence *et al*., 2016). This increase drives the expansion of the maturing spine head and coincides with shortening of the filopodia precursor and constriction at the base of the maturing spine. These changes yield the prototypical mature, mushroom-shaped spine possessing an enlarged head filled with branched actin that is connected to the dendrite by a narrow spine neck containing both branched and unbranched actin filaments.

Guanine nucleotide exchange factors (GEFs) for Rho-related GTPases likely play key roles in the regulation of actin dynamics underlying both spine formation and spine structural plasticity, as these proteins link signal transduction pathways to the activation of factors required for actin assembly and turnover (Newey *et al*., 2005; Kiraly *et al*., 2010; Chen, Wirth and Ponimaskin, 2012; Penzes and Cahill, 2012; Tejada-Simon, 2015; Woolfrey and Srivastava, 2016). One GEF that has been implicated repeatedly in driving spine maturation is the Rac1 and Cdc42 GEF β-Pix (Zhang *et al*., 2003, 2005; Za *et al*., 2006; Penzes *et al*., 2008; Saneyoshi *et al*., 2008; Kiraly *et al*., 2010). The activation of Rac1 and Cdc42 within spines by β-Pix is thought to drive spine maturation via several distinct pathways. First, GTP-bound Rac1 and GTP-bound Cdc42 can activate two nucleation promoting factors (NPFs) for the Arp2/3 complex (WAVE in the case of Rac1-GTP, WASp in the case of Cdc42-GTP), resulting in the activation of branched actin filament assembly (Rotty, Wu and Bear, 2013). Several published studies are consistent with the idea that β-Pix promotes spine maturation via this pathway (Kim *et al*., 2006; Soderling *et al*., 2007; Rácz and Weinberg, 2008; Wegner *et al*., 2008; Chen, Wirth and Ponimaskin, 2012; Tejada-Simon, 2015; Spence *et al*., 2016). Second, both GTP-bound Rac1 and GTP-bound Cdc42 can activate PAK kinase, leading to the activation of myosin 2 filament assembly and contractility within spines via the PAK-dependent phosphorylation of the myosin’s regulatory light chain (RLC) (Beach and Hammer, 2015). Direct evidence that this β-Pix-dependent pathway promotes spine maturation has been presented (Zhang *et al*., 2003, 2005). Active PAK also appears to promote spine maturation by phosphorylating/activating LIM-kinase (LIM-K), as this leads to the stabilization of spine actin via the LIM-K-dependent inhibition cofilin-driven actin filament turnover (Meng *et al*., 2002; Chen *et al*., 2007). Finally, the morphogenesis of spines that occurs in response to NMDA receptor activation has been shown to be mediated by a signaling complex containing two related calcium-calmodulin-dependent protein kinases, β-Pix, and Rac1 (Penzes *et al*., 2008; Saneyoshi *et al*., 2008).

In most cellular contexts where F-actin plays a critical role, myosin 2 also plays a critical role. Consistent with this paradigm, myosin 2B is enriched along with F-actin in spines, and attenuating the myosin’s function in developing neurons either pharmacologically or by knockdown results in a strong block in spine maturation (Morales and Fifková, 1989; Ma *et al*., 2006; Ryu *et al*., 2006; Rex *et al*., 2010; Hodges *et al*., 2011; Rubio *et al*., 2011; Gavin *et al*., 2012; Koskinen *et al*., 2014; Ozkan *et al*., 2015; Briggs *et al*., 2018). Moreover, blocking myosin 2B function in fully-developed neurons shows that it also plays a critical role in the activity-dependent changes in spine actin morphology underlying learning and memory (Rex *et al*., 2010; Rubio *et al*., 2011; Gavin *et al*., 2012; Ozkan *et al*., 2015; Briggs *et al*., 2018). While myosin 2B is present in dendritic filopodia as well as in mature spines (Korobova and Svitkina, 2010), its function in mature spines has been studied most. Myosin 2B resides primarily at the base of these structures (i.e. in the spine neck and base of the head), most likely in the form of bipolar filaments (Korobova and Svitkina, 2010). The contractile activity of these bipolar filaments is thought to drive the shortening of immature spines during spine maturation, the constriction of the spine neck, the tip-to-base flow of a dynamic spine actin pool, the cross-linking of a stable spine actin pool, and possibly global spine actin dynamics by catalyzing actin filament turnover downstream of myosin-dependent actin filament breakage (Ozkan *et al*., 2015). Interestingly, Purkinje neurons express an alternatively spliced version of myosin 2B known as myosin 2B-B2 in addition to regular myosin 2B (Ma *et al*., 2006). Mice lacking this splice variant exhibit aberrant Purkinje neuron development (reduced numbers of dendritic spines and branches) and impaired motor coordination (Ma *et al*., 2006).

The focus of this study is on the function of the myosin 2-like protein myosin 18A (Billington *et al*., 2015; Horsthemke *et al*., 2019). Like myosin 2, myosin 18A possesses a globular head domain followed by a rod-like coiled-coil tail domain. Unlike myosin 2, myosin 18A also possesses N- and C-terminal extensions that harbor both recognizable (e.g. PDZ domain, SH3 domain binding sites) and uncharacterized protein: protein interaction domains (Billington *et al*., 2015; Horsthemke *et al*., 2019). Also unlike myosin 2, myosin 18A does not exhibit motor activity and does not self-assemble into bipolar filaments, the working form of myosin 2 (Guzik-Lendrum *et al*., 2011, 2013). Despite these peculiarities, and the fact that myosin 18A is present in cells in amounts that are significantly sub-stochiometric to myosin 2 (about 1 myosin 18A for every 10 to 50 myosin 2s) (Billington *et al*., 2015), the knockout of myosin 18A results in embryonic lethality in both flies (Bonn *et al*., 2013) and mice (Horsthemke *et al*., 2019). Moreover, at least one spliced isoform of myosin 18A appears to be expressed in every differentiated mammalian cell type based on expression profiling.

A major breakthrough in resolving the enigma of myosin 18A function came from data showing that purified myosin 18A co-assembles with purified myosin 2 *in vitro* to make mixed filaments, and that such mixed filaments can be seen in living cells using super-resolution light microscopy (Billington *et al*., 2015). Importantly, co-assembly provided explanations for myosin 18A’s unusual properties. First, it does not need to self-assemble because it can co-assemble with myosin 2. Second, it does not need to be a motor because myosin 2 in the mixed filament will provide that function. Third, its low stoichiometry relative to myosin 2 should result in mixed filaments containing mostly myosin 2 with a few myosin 18A molecules sprinkled in. Finally, co-assembly suggested a clear role for myosin 18A: display of its N- and C-terminal protein interaction domains on the surface of mixed filaments might serve to recruit proteins to mixed filaments and/or attach them to cellular structures. Implicit in this hypothesis is that myosin 18A function is inextricably linked to myosin 2 function. Given that myosin 18A is ubiquitously expressed, it may be the case that most past studies of myosin 2 function in organisms ranging from fly to man were actually interrogating the function of mixed filaments of myosin 2 and myosin 18A.

One bona fide myosin 18A binding partner is β-Pix, which binds via 12 residues near its C-terminus to myosin 18A’s ∼80-residue non-helical tailpiece (Hsu *et al*., 2010, 2014). This interaction has been shown to regulate cell migration and focal adhesion dynamics in mesenchymal cells (Hsu *et al*., 2010, 2014). A second binding partner is the myosin 2 regulatory light chain kinase MRCK, which binds to myosin 18A’s N-terminal PDZ domain through the adaptor protein LRAP (Tan *et al*., 2008). This interaction has been shown to mediate myosin 18A’s role in regulating the dynamics of actomyosin 2 networks in mesenchymal cells by phosphorylating and activating myosin 2 (Tan *et al*., 2008). Relevant to this, β-Pix is typically found in a complex with GIT1 and PAK kinase, so the interaction of myosin 18A with β-Pix could conceivably lead to myosin 2 activation downstream of PAK (Zhang *et al*., 2005; Za *et al*., 2006).

Given that both myosin 2 and β-Pix play important roles in spine maturation and function, that myosin 18A co-assembles with myosin 2 and binds β-Pix, and that myosin 18A is highly expressed throughout the brain (e.g. in pyramidal neurons in the hippocampus and cortex, in cerebellar Purkinje neurons), we decided to investigate myosin 18A function in a neuronal cell type. With regard to which cell type to study, previous work from our lab showed that myosin Va transports tubules of smooth endoplasmic reticulum into Purkinje neuron spines to support synaptic plasticity and motor learning, thereby explaining the ataxia exhibited by *dilute*/myosin Va null mice (Wagner, Brenowitz and Hammer, 2011). Our subsequent efforts have resulted in improvements in culturing and transfecting Purkinje neurons, in the development of a Purkinje neuron-specific miRNA-mediated knockdown system (Alexander and Hammer, 2016), and in creating Purkinje neurons from embryonic stem cells (Alexander and Hammer, 2019). For these and other reasons we decided to investigate myosin 18A function in Purkinje neurons. Here we show that myosin 18A targets β-Pix to Purkinje neuron spines to promote spine maturation.

## METHODS

### Mice and primary cerebellar cultures

Timed-pregnant C57BL/6 females of gestation day 17-18 were purchased from Charles Rivers Laboratories. E17-E18 embryos were harvested and mixed cerebellar cultures were prepared as described previously (Alexander and Hammer, 2016). Animals were anesthetized by inhalation of Isoflurane and euthanized by cervical dislocation. The generation of the myosin 18A^flox/flox^ conditional knockout (cKO) mouse will be described in a separate publication. Efforts were made to limit the number of animals used in this study. All animal experiments were approved by the Institutional Animal Care and Use Committee of the National Heart, Lung, and Blood Institute in accordance with the National Institutes of Health guidelines.

### Immunofluorescence

Primary cerebellar cultures plated on 35mm round dishes with coverslip bottoms were fixed for 15 minutes at room temperature in 4% paraformaldehyde (PFA) in Cytoskeleton Fixation Buffer (10 mM PIPES (pH 6.8), 100 mM NaCl, 10% (v/v) sucrose, 3 mM MgCl_2_, 1 mM EGTA) by diluting a 16% PFA stock solution (Electron Microscope Sciences, 15710). Samples were then washed three times with PBS (Gibco, 70011044), permeabilized in PERM buffer (300 mM Glycine, 0.5% Triton X-100 in PBS (pH 7.4)) for 10 minutes, washed once with PBS, and blocked for 30 minutes in IF Blocking Buffer (3% (w/v) BSA (Sigma A8412), 0.1 % (v/v) Tween 20 in PBS (pH 7.4)). Purkinje neurons were identified using anti-Calbindin D28K antibody (guinea pig, 1:500, Synaptic Systems, 214 004) or anti-mCherry antibody (mouse, 1:500, MyBioSource, MBS555186). Depending on the experiment, cultures were also stained with the following primary antibodies at the indicated dilutions: an anti-myosin 18Aα raised against the C-terminal 18 residues of myosin 18Aα/β (rabbit, 1:50) (Billington *et al*., 2015; Bruun *et al*., 2017), anti-Bassoon (rabbit, 1:200, Synaptic Systems, 141 003), anti-myosin 2B (rabbit, 1:200, ThermoFisher, PA517026), or anti-β-Pix (mouse, 1:200, BD Biosciences, 611648) in 200 µl of IF Blocking Solution for 45 minutes at room temperature. Following three 5-minute washes in PBS, samples were incubated for 45 minutes at room temperature with 1:500 dilutions of the appropriate labeled secondary antibody (Alexa Fluor 546 goat anti-guinea pig, Thermo Fisher, A11074; Alexa Fluor 546 goat anti-mouse, Thermo Fisher, A11003; or Alexa Fluor 488 goat anti-rabbit, Thermo Fisher, A11008). Samples were then washed three times with PBS. To visualize F-actin, cells were incubated during the final wash step with Alexa Fluor 647 labelled Phalloidin for 5 minutes (1:100, Thermo Fisher Scientific, A12379). Following all wash steps, cells were overlaid with 200 μl of Fluromount-G mounting medium (Electron Microscope Sciences, 179840-25), covered with a 25 mm circular coverslip (Electron Microscope Sciences, 72223-01), and sealed with slide sealant. Images were obtained using a Zeiss LSM 780 laser scanning confocal microscope equipped with a Zeiss 63 X, 1.4 NA oil objective. Perfusion fixed sagittal cerebellar slices were a kind gift of Dr. Herbert Geller (NHLBI/NIH) and prepared as previously described (Yi *et al*., 2012).

### Cloning

All oligonucleotides used in this study were purchased from Eurofins Genomics at 50 nM scale and purified by standard desalting. Table 1 shows the oligonucleotides used to generate the pL7-based (Wagner, McCroskery and Hammer, 2011), Purkinje neuron-specific expression plasmids for myosin 18Aα, myosin 18Aβ, myosin 2B, and β-Pix using the following DNA templates: mouse myosin 18A (Riken clone F730020C19), mouse myosin 2B-B2 (Ma *et al*., 2006), and mouse β-Pix (Mammalian Gene Collection clone MMM1013-202702676). DNA fragments containing 21 base pair overhangs homologous to the vector backbone were generated by PCR using Phusion High-Fidelity DNA polymerase (NEB, M0530), purified, and inserted into Bgl II/Sal I-linearized pL7 mGFP or pL7 mCherry (Wagner, McCroskery and Hammer, 2011) using an In-Fusion cloning kit (Takara, 638920). The specific fragments generated using the appropriate template and indicated primers were: myosin 18Aα (oM18_1 and oM18_2), myosin 18Aβ (oM18_3 and oM18_4), myosin 2B-B2 (oM18_5 and oM18_6), myosin 18Aα-ΔNHT (lacking C-terminal residues 1940-2050) (oM18_7 and oM18_8), myosin 18Aα/β-CC (containing residues 1241-2035 of myosin 18Aα, which are common to myosin 18Aα and myosin 18Aβ) (oM18_9 and oM18_10), and β-Pix (oM18_19 and oM18_20). pL7 mGFP myosin 18Aα-AB^mut^ in which residues V117 and L118 were both changed to A (Taft *et al*., 2013) was generated by site directed mutagenesis using primers oM18_17 and oM18_18 (Table 1) and a Quikchange II Site-directed Mutagenesis kit (Agilent, 200523). The PCR product was digested with DpnI enzyme to remove template vector and transformed into Stbl2 chemically competent cells (Thermo Fisher, 10268019). β-Pix lacking its myosin 18A binding site (residues 639-647) was synthesized by Gene Universal and cloned into pL7 mGFP as a Bgl II/Sal I fragment. The myosin 18Aα N-terminal (NT) constructs were generated using traditional cloning methods. Specifically, the wild type N-terminus (residues 1-334) (NT), the N-terminus lacking the KE-rich region (residues 1-29 deleted) (NT-ΔKE), the N-terminus containing the PDZ domain mutation (residue G232 changed to P; (Tonikian *et al*., 2008)) (NT-PDZ^mut^), and the N-terminus containing the actin binding site mutation (NT-AB^mut^) were amplified by PCR with Bgl II/Sal I ends using the appropriate template and primers indicated in Table 1, or they were synthesized by Gene Universal. The purified NT fragments were digested with Bgl II/Sal I and ligated into Bgl II/Sal I-linearized pL7 mGFP. To generate the *E. coli* expression plasmids NT (residues 1-334) and NT-AB^mut^ (containing the VLΔAA point mutations), fragments with Nco I/Sal I ends were PCR amplified using the appropriate pL7 template and primers oM18_21 and oM18_22. The PCR products were purified, digested with Nco I/Sal I, and ligated into the GST expression plasmid pGEX-Parallel 2 (Sheffield, Garrard and Derewenda, 1999) linearized with Nco I/Sal I to generate GST fusion proteins NT and NT-AB^mut^. All plasmids generated in this study were confirmed by sequencing.

**Table 1.**
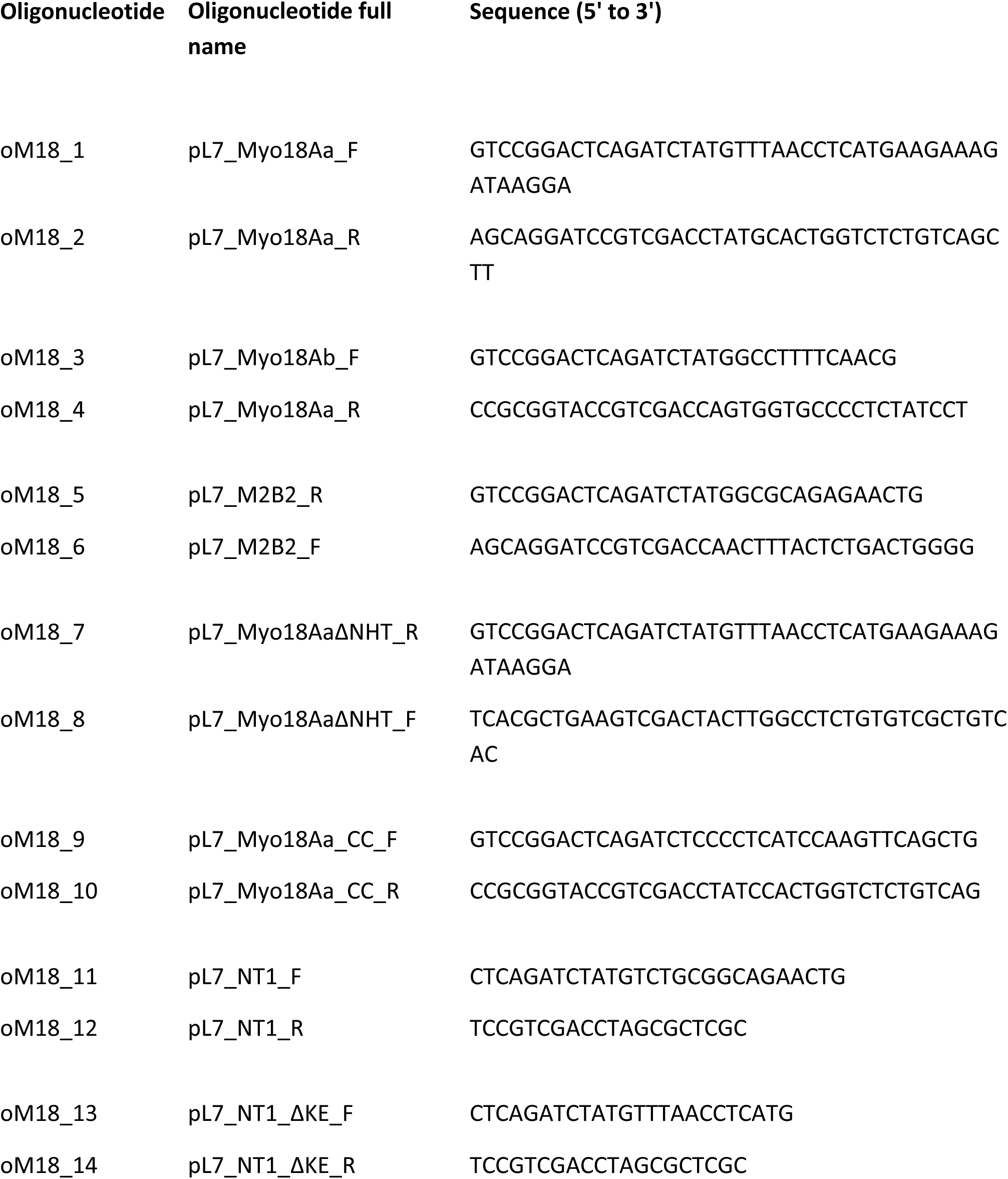
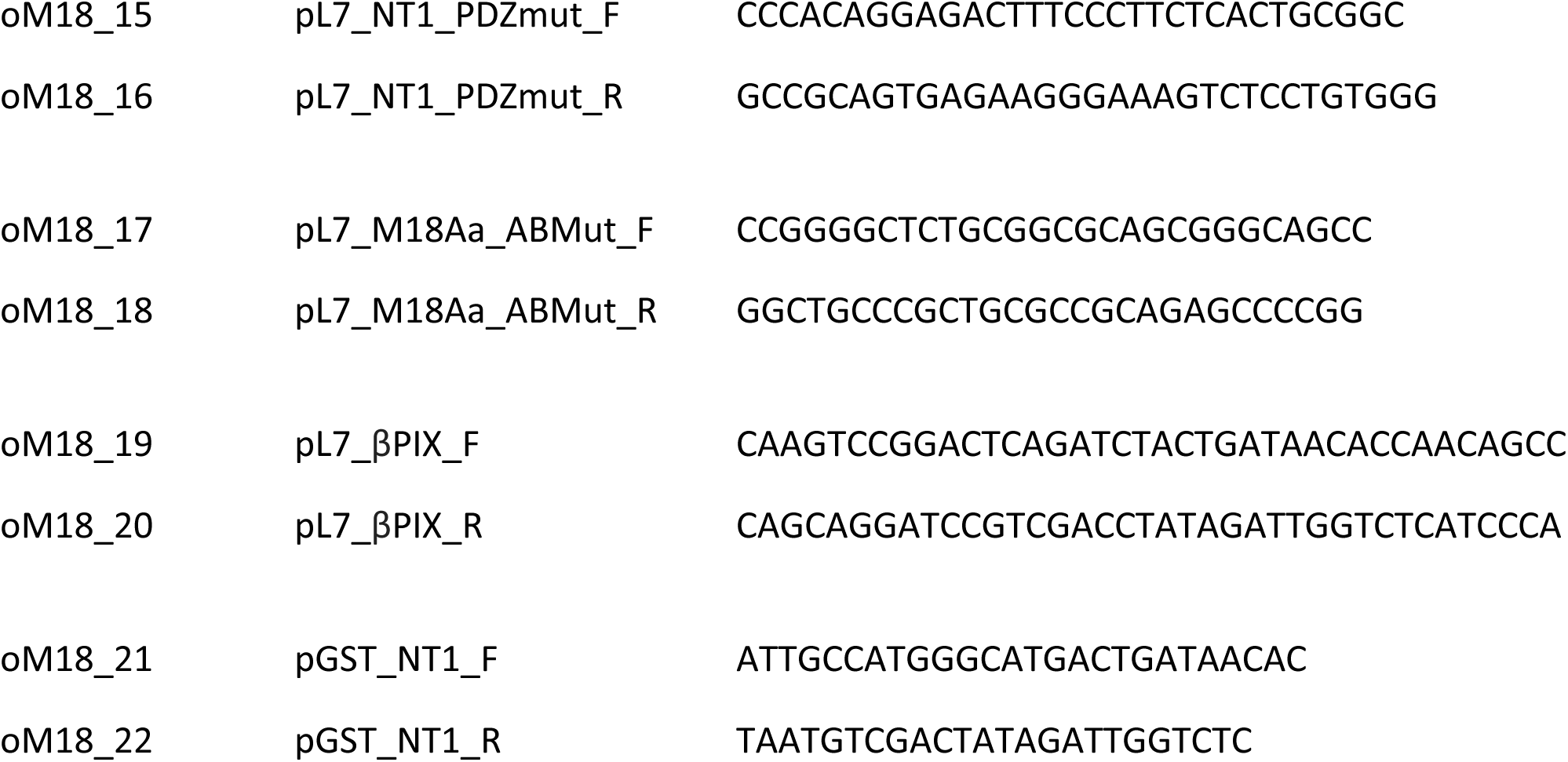
Oligonucleotides used in this study.

### miRNA knockdown

Purkinje neuron-specific, miRNA-mediated protein knockdown was performed as described previously (Alexander and Hammer, 2016) using the BLOCK-iT Inducible Pol II miR RNAi Expression Vector Kit (Life Technologies K4939-00). Briefly, complementary oligonucleotides for generating miRNAs directed against the 3’ UTR of target proteins were designed using the Thermo Fisher BLOCK-iT RNAi Design Tool (https://rnaidesigner.thermofisher.com/rnaexpress/) and NM_001291213 (mouse myosin 18A) and NM_001346804 (mouse β-PIX) as reference sequences. The oligonucleotides were ligated into the pcDNA6.2/emGFP plasmid provided with the kit and the resulting plasmids used as templates to PCR amplify complete miRNA cassettes with Bgl II and Sal I ends for ligation into Bgl II/Sal I cut pL7 mCherry (Alexander and Hammer, 2016). The final constructs were sequence verified.

### Western Blotting

Cerebella from E17 embryos and adult (∼8 weeks old) mice were harvested and lysed in RIPA buffer (50 mM Tris (pH 7.4), 150 mM NaCl, 2 mM EDTA, 0.5% (w/v) Na-deocycholate, 1% (v/v) NP-40, and 0.1% (v/v) SDS) supplemented with Complete Protease Inhibitor Cocktail (Roche, 11697498001)). Whole cell extracts of Jurkat cells, a human leukemic T cell line (ATCC, TIB-152), were prepared as described previously (Murugesan *et al*., 2016). Protein concentrations were determined using a BCA Protein Assay (Thermo Fisher, 23225). Equal amounts of protein were run on 6% SDS-PAGE gels and blotted onto nitrocellulose using a semi-dry Trans-Blot (Bio-Rad, 1703940). Blots were incubated overnight at 4°C with a rabbit anti-myosin 18A polyclonal antibody that recognizes the C-terminal 18 residues of both myosin 18Aα and 18Aβ (Bruun *et al*., 2017) diluted 1:1000 in TRIS-buffered saline containing 0.05% (v/v) Tween-20 (TBS-T) and supplemented with 5% (w/v) dry nonfat milk. Blots were washed extensively with TBS-T and then incubated with anti-rabbit HRP secondary antibody (GE Lifesciences, NA934) diluted 1:5000 in TBS-T supplemented with 5% (w/v) dry nonfat milk. Bands were detected using the SuperSignal West Pico PLUS Chemiluminescent detection system (Thermo Fisher, 34577). The blot in Figure S1 also contained an extract of adult mouse cardiac muscle, and was probed with a rabbit anti-myosin 18A polyclonal antibody raised against the C-terminal 300 residues of myosin 18Aα and myosin 18β (Bruun *et al*., 2017) (this antibody should also see myosin 18Aγ because it contains the N-terminal 252 residues of the immunogen). The extent of miRNA-mediated knockdown of myosin 18A was estimated using NIH 3T3 mouse embryonic fibroblasts (ATCC CRL-1658) cultured in DMEM medium containing 4 mM Glutamax (Thermo Fisher, 10566) and 10% (v/v) FBS. Cells were grown to ∼70% confluency, harvested with TripLE dissociation reagent (Thermo Fisher, 12604013) and counted. Each miRNA, under the control of the CMV promoter, was introduced into cells by nucleofection using an Amaxa Nucleofector (Lonza). Briefly, 1 x 10^6^ cells were collected and resuspended in 100 ul of Nucleofection Solution (5 mM KCl, 15 mM MgCl_2_, 120 mM Na_2_HPO_4_/NaH_2_PO_4_ (pH 7.2), 50 mM Mannitol) containing 2 ug of plasmid DNA. Nucleofection was carried out using program U-(0)03. Cells were immediately transferred to one well of a 6-well plate containing 3 ml of growth medium, cultured for 24 h, harvested, lysed in SDS-PAGE loading buffer, and subjected to SDS-PAGE. Proteins were transferred to nitrocellulose, probed with the rabbit anti-myosin 18A polyclonal antibody directed against the C-terminal 18 residues of myosin 18Aα/β (1:1000, cite JB Cytoskeleton 2017) and with a mouse anti-RACK1 monoclonal antibody (1:1000, Santa Cruz, sc-17754) as the loading control, and developed using an anti-rabbit HRP secondary antibody (1:5000, GE Lifesciences, NA931) and an anti-mouse HRP secondary antibody (1:10,000, GE Healthcare, NA935). Blots were imaged using an Amersham Imager 600 (GE Healthcare Life Sciences) and the extent of knockdown was estimated by densitometry using Fiji (Rueden *et al*., 2017). For the Western blot in Figure S5, Panel D, mouse embryo fibroblasts (MEFs) from the myosin 18A cKO mouse were prepared and cultured as described previously (Beach *et al*., 2017), transfected with a puromycin resistance plasmid that expresses Cre recombinase (Addgene, 17408), grown for 48 hours and then selected for using 2.5 μg/ml puromycin (Invivogen, ant-pr), and processed as for the miRNA-treated 3T3 fibroblasts except that a rabbit polyclonal against the heavy chain of clathrin (1:1000, Abcam, ab21679) was used as the loading control.

### F-actin co-sedimentation assay

GST-NT and GST-NT-AB^mut^ were expressed in BL21-CodonPlus (DE3)-RIPL cells (Agilent, 230280), purified as described previously (Fujiwara *et al*., 2014), and dialyzed into KMEI buffer (50 mM KCl, 1 mM MgCl_2_, 1 mM EGTA and 10 mM imidazole (pH 7.0)). Actin was purified as described previously (Fujiwara, Remmert and Hammer, 2010) and dialyzed into G-buffer (2 mM TRIS (pH 8.0), 0.2 mM ATP, 0.5 mM DTT, 0.1 mM CaCl_2_ and 1 mM NaN_3_). The affinities of NT and NT-AB^mut^ for F-actin were determined by high speed co-sedimentation assays essentially as described previously (Fujiwara, Remmert and Hammer, 2010; Fujiwara *et al*., 2014). Briefly, 1 μM of NT or NT-AB^mut^ were incubated for 60 min at room temperature with increasing molar ratios of F-actin (1:0, 1:1, 1:2.5, 1:3.5 or 1:5 for NT; 1:0, 1:10, 1:15, 1:25 or 1:40 for NT-AB^mut^), centrifuged at 100,000 x *g* for 1 h, and the supernatant (S) and pellet (P) fractions resolved by SDS-PAGE. The amount of NT fusion protein in each sample was determined by densitometry using Fiji (Rueden *et al*., 2017). The data was fit to a one-site binding hyperbola in obtain K_d_ values.

### Image analyses

Stained cells were imaged using a Zeiss LSM 780 confocal microscope equipped with 63 X Plan Apo objectives (NA 1.4). All morphological measurements of Purkinje neurons were performed using Fiji Image Software (Rueden *et al*., 2017). To measure dendritic spine length, serial 0.4 μm confocal sections that encompass the entire spine (identified using the cell volume marker mCherry in the 546 nm channel) together with the adjacent dendrite were summed using the Fiji Z projection function. Using the line tool, individual dendritic spines were masked by drawing a line from the tip of the spine head through to the dendrite at the base of the spine. Spine length was then determined by measuring the length of each corresponding line. To determine dendritic spine density, four serial 0.4 μm confocal sections (a total of 1.2 μm) were summed using the Fiji Z projection function. The number of dendritic spines in 10 μm lengths of dendrite were counted and used to determine the average number of spines per μm^2^. The enrichment of proteins in dendritic spines relative to dendrites was determined by ratio imaging as described previously (Alexander *et al*., 2018). Briefly, cells were co-transfected with a GFP-tagged protein of interest and mCherry as a volume marker. Serial 0.4 μm confocal sections that encompass an entire spine (identified using the mCherry volume marker in the 546 nm channel) together with the adjacent dendrite were summed using the Fiji Z projection function. The total fluorescence for the GFP signal (488 nm channel) in the spine and in an equal-sized area of the dendrite immediately under the spine were then measured. To correct for differences in volume between these two regions, the total fluorescence for the mCherry volume marker (546 nm channel) was also measured in both regions. The apparent fold-enrichment of the GFP-tagged protein in the spine relative to the subjacent dendrite region was then corrected for any difference in volume between these two regions to obtain an accurate value for the fold-enrichment of the GFP-tagged protein in the spine. To determine the content of F-actin per spine, cells were fixed and stained with a rabbit anti-mCherry antibody (1:200, MyBioSource, MBS9400754) followed by an Alexa 546-conjugated goat anti-rabbit secondary antibody (1:500, Thermo Fisher, A11035) (this staining serves to both identify myosin 18Aα miRNA-expressing cells and mark their volume), and with Alexa 647-conjugated Phalloidin to label F-actin. Control cells were fixed and stained with Alexa 647-conjugated Phalloidin and with anti-Calbindin D28K followed by Alexa 546-conjugated anti-Guinea Pig secondary antibody to mark their volume. Serial 0.4 μm confocal sections that encompass the entire dendrite were collected and summed using the Fiji Z projection function. The mCherry or anti-Calbindin volume signals were then thresholded to determine the outline of the dendrite and dendritic spines, and this outline was then used to mask any Phalloidin signal in the 647 channel outside the Purkinje neuron volume. The total Phalloidin fluorescence per spine was then measured. To determine the content of myosin 2B per spine, cells were fixed and stained with the anti-mCherry antibody as described above to identify cells expressing the myosin 18Aα or Scrambled miRNA and to mark their volume, and with the rabbit anti-myosin 2B antibody (1:200, ThermoFisher, PA517026) to label the myosin. The content of myosin 2B per spine was determined as for F-actin. As discussed in the relevant Figure legends, we also provided estimates of the reductions in spine F-actin and myosin 2B content in knockdown cells that included corrections for differences between control and knockdown cells in spine volume.

### Statistical Analyses

Statistics for all experiments were performed using Graphpad Prism v7 software (Mac Version), and all figures were prepared using Inkscape, Inkscape Project v0.92.3 (Mac Version). Statistical details, including N values and what these values represent, are included in the appropriate figures, figure legends and text. For comparison of two groups, a parametric Student’s *t*-test with Welch’s correction was performed. For the Pearson’s correlation coefficient data in Figure S2, the signals in the 594 channel (Calbindin D28K) and 488 channel (myosin 18Aα) were measured using the Fiji co-localization tool. As a control, the 488 channel (myosin 18Aα) was rotated 90 degrees and the Pearson’s correlation coefficient was re-calculated. Three equal-sized regions of cerebellar slices were analyzed. All of the statistical data is presented as means with standard deviations. Significance values, which are included in figures, figure legends, and text, are defined as following: * *p*=<0.05, ** *p*=<0.01, *** *p*=<0.001, **** *p*=<0.0001.

### Immunoelectron microscopy

Immunoelectron microscopy was performed using a pre-embedding immunoperoxidase method followed by silver/gold enhancement as described previously by Petralia *et al*. (2010) and He *et al*. (2014). Briefly, two 5-week old Sprague-Dawley rats were anesthetized with Ketamine/xylazine and perfused with 4% paraformaldehyde in PBS. Fifty micrometer brain sections in 30% sucrose/PBS were frozen using acetone in dry ice and stored at −80°C. Thawed sections were washed in PBS and incubated for 1 hour in 10% (v/v) normal goat serum diluted in PBS, and then in a 1:2000 dilution of the rabbit C-terminal myosin 18Aα primary antibody in PBS at 4°C overnight. Samples were processed the following day using a Vectastain kit (biotinylated goat-anti-rabbit igG secondary antibody and avidin-biotin-peroxidase complex) and ImpacDAB (Vector Laboratories). Control sections lacking the primary antibody were unlabeled. Subsequent silver/gold toning was performed in cacodylate buffer, including incubation for 10 minutes at 60°C in 0.2% (w/v) silver nitrate, 0.2% (w/v) sodium borate and 2.6% (v/v) hexamethylenetetramine, followed by 2 minutes each with 0.05% (w/v) gold chloride and 3% (w/v) sodium thiosulfate. Finally, sections were fixed in 2% (w/v) glutaraldehyde and then 1% (w/v) osmium tetroxide, dehydrated in alcohols and propylene oxide, and embedded in Epon. Thin sections from the 2 rats were produced on a Leica Ultracut ultramicrotome and were examined in a JEOL JSM-1010 transmission electron microscope.

## RESULTS

### Purkinje neurons express myosin 18Aα

As discussed briefly in the Introduction, myosin 18A mRNA (detected using a probe that should recognize all spliced isoforms) is present at moderate to high levels throughout the central nervous system, including in cerebellar Purkinje neurons (Lein *et al*., 2007). Three myosin 18A splice isoforms have been described to date: 18Aα, 18Aβ, and 18Aγ (Billington *et al*., 2015; Horsthemke *et al*., 2019). These three isoforms share head and coiled-coil domains but differ in their N- and C-terminal extensions, with myosin 18Aβ lacking myosin 18Aα’s N-terminal extension, and myosin 18Aγ having large, unique N- and C-terminal domains (Figure 1A). A Western blot of embryonic and adult mouse cerebellar extracts (Figure 1B, lanes 2 and 3, respectively) probed with an antibody that recognizes the C-terminal 18 residues of both myosin 18Aα and 18Aβ (Bruun *et al*., 2017) detected only the ∼235 kDa myosin 18Aα isoform in both cerebellar extracts (lane 1 shows the strong signal for myosin 18Aβ at ∼200 kDa in a Jurkat T cell whole cell extract). A second Western blot of extracts from Jurkat T cells, adult mouse cerebellum and adult mouse cardiac muscle probed with an antibody to the C-terminal 300 residues of myosin 18Aα/myosin 18Aβ, which overlap with 252 residues within the tail domain of myosin 18Ay, again detected only myosin 18Aβ in T cells and myosin 18Aα in the cerebellar extract (Figure S1, lanes 1 and 2, respectively), but also a ∼290 kDa isoform specific to the cardiac muscle extract (Figure S1, lane 3). This ∼290 kDa band probably corresponds to myosin 18Aƴ given that its expression is reported to be restricted to cardiac muscle (Horsthemke *et al*., 2019); our RT PCR data also shows that myosin 18Aƴ mRNA is not detectable in adult mouse brain (data not shown)). Based on all of these observations, we conclude that Purkinje neurons express myosin 18A, and that they express predominantly if not exclusively the 18Aα isoform.

**Figure 1.**
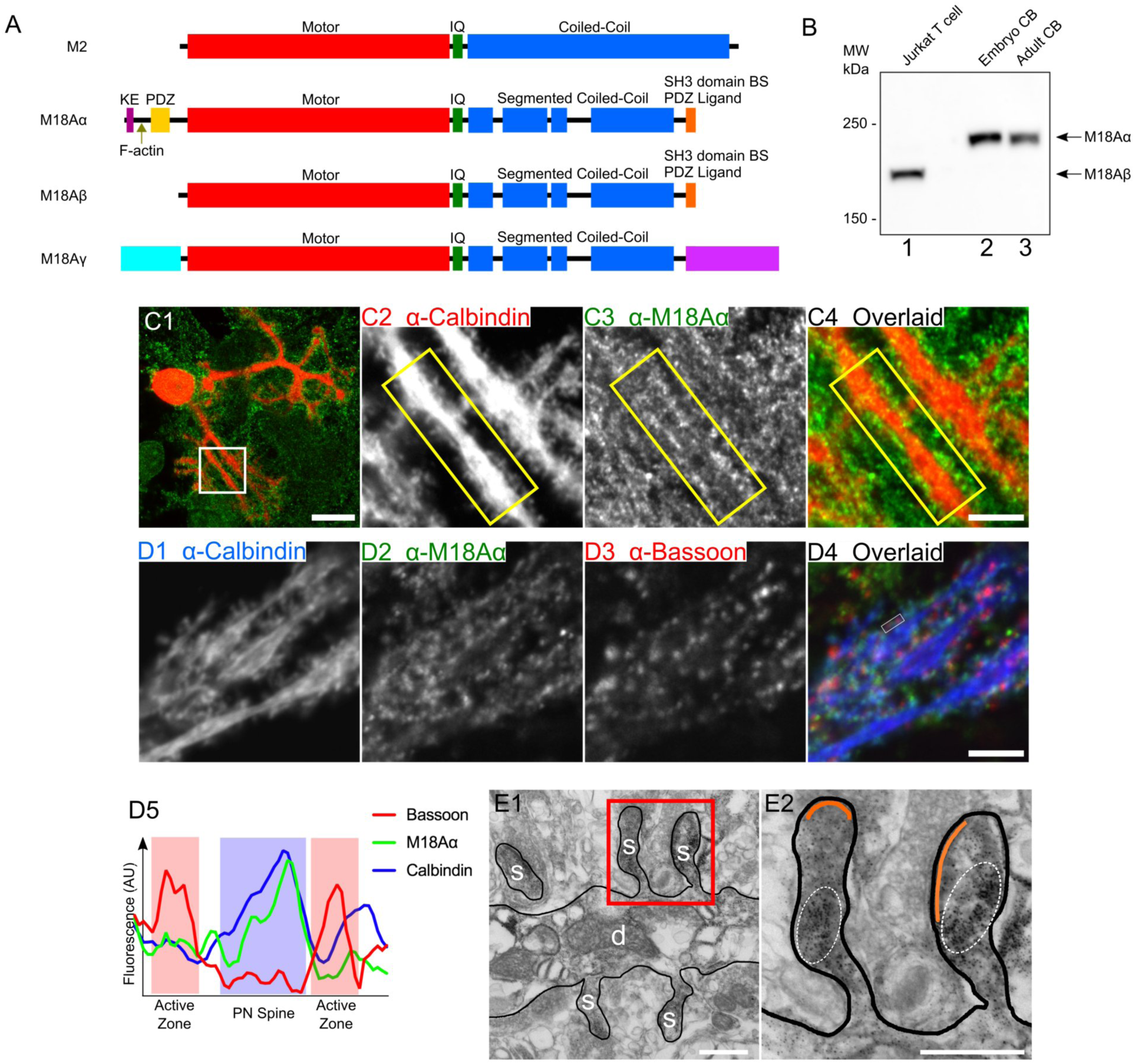
Endogenous myosin 18Aα localizes to the dendritic spines of Purkinje neurons. (A) Domain architecture for myosin 2 (M2) and the three known isoforms of myosin 18A, myosin 18Aα (M18Aα), myosin 18Aβ (M18Aβ), and myosin 18Aγ (M18Aγ). Domains discussed in the text are labeled. (B) Representative Western blot performed on a human Jurkat T cell extract (lane 1), a whole mouse embryonic cerebellum extract (lane 2), and a whole mouse adult cerebellum extract (lane 3), and probed with an anti-myosin 18A antibody against the C-terminal 18 residues of myosin 18Aα/myosin 18Aβ. The expected positions for the heavy chains of myosin 18Aα and myosin 18Aβ are indicated. (C1-C4) Panel C1 shows a representative cultured Purkinje neuron (DIV 18) that was double-stained for Calbindin D28K in red and myosin 18Aα in green, while Panels C2-C4 show enlargements of the boxed region in Panel C1. The yellow boxes highlight the localization of endogenous myosin 18Aα in dendritic spines. (D1-D5) Panels D1-D4 show a portion of a dendrite from a cultured Purkinje neuron (DIV 18) that was tripled-stained for Calbindin D28K in blue (D1), myosin 18Aα in green (D2), and the granule neuron pre-synaptic marker Bassoon in red (D3) (D4, Overlaid image), while Panel D5 shows a line scan across a representative spine bounded by closely-associated Bassoon signals (Calbindin signal, blue line; myosin 18Aα signal, green line; Bassoon signal, red line; Purkinje neuron spine, blue shading; presynaptic active zones, red shading). (E1 and E2) Panel E1 shows a representative electron micrograph taken of a cerebellar section from a five-week old Sprague-Dawley rat that was perfusion-fixed and labeled with a pre-embedding immunoperoxidase method for myosin 18Aα, followed by silver/gold enhancement. The dendrite (d) and several spines (s) are labeled. Panel E2 shows an enlargement of the two spines present within the boxed region in Panel E1. The area within each spine that is enriched for myosin 18Aα immunogold labeling (black dots) is encircled by a dashed line, while the PSD in each spine is pseudo-colored orange. The image in E1 is representative of over 200 images taken of several sections obtained from two rats. Scale bars: 20 μm (C1 and D1), 5 μm (C4 and D5), and 500 nm (E1 and E2).

### Endogenous myosin 18Aα localizes to Purkinje neuron spines

To demonstrate more directly that Purkinje neurons express myosin 18Aα, and to begin to define its intracellular localization, we stained primary cerebellar cultures with an antibody to the Purkinje neuron-specific marker Calbindin D28K in red to identify Purkinje neurons (this staining also acts as a fill marker to reveal dendritic spines) (Yang *et al*., 2000; Laure-Kamionowska and Málińska, 2009; Wagner, McCroskery and Hammer, 2011; Alexander *et al*., 2018; Alexander and Hammer, 2019), and with the antibody to the C-terminal 18 residues of myosin 18Aα in green. Figure 1, Panel C1, shows a representative stained Purkinje neuron, while Panels C2 through C4 show enlarged images of the individual channels and the overlaid image for the boxed region in Panel C1. Careful inspection reveals that the edges of the dendrites present within this boxed region are studded with Calbindin-positive spines that contain the majority of the signal for endogenous myosin 18Aα (compare the positions of the two signals within the yellow boxes in Panels C2 through C4). Similar results were obtained using frozen sections of cerebellar tissue that were fixed and stained for Calbindin and myosin 18Aα (Figure S2; see the arrowheads). Together, these initial observations argue that Purkinje neurons do indeed express myosin 18Aα, and that this myosin localizes primarily to dendritic spines.

Cultured Purkinje neurons make extensive synaptic contacts with granule neurons that are also present in primary cultures (Richter *et al*., 1999; Seil, 2014; Alexander *et al*., 2018). To address the possibility that the spine-associated signals described above might actually be on the presynaptic side, we stained primary Purkinje neuron cultures for Calbindin in blue, myosin 18Aα in green, and the presynaptic marker Bassoon in red (see Panels D1-D3, respectively, in Figure 1). Inspection of the overlaid image in Panel D4 shows that while the green signals for myosin 18Aα and the red signals for Bassoon are clearly quite close, they appear to be largely non-overlapping. Line scans bore this out, as shown for a representative spine in Panel D5, where the signals for myosin 18Aα (green line) and Calbindin (blue line) marking the spine (blue shading), and the signal for Bassoon (red line) marking two adjacent presynaptic active zones (red shading), are largely non-overlapping. To provide additional evidence that myosin 18Aα is postsynaptic, rat cerebellar tissue that had been perfusion-fixed was labeled with a pre-embedding immunoperoxidase method using the C-terminal myosin 18Aα antibody, followed by silver/gold enhancement. Figure 1, Panel E1, and the enlarged inset in Panel E2, show that the myosin 18Aα signal, represented by the mottled black precipitate encircled with dashed lines, lies within the Purkinje neuron spine and well below the post synaptic density (PSD, pseudo-colored in orange). Together, these observations indicate that myosin 18Aα is expressed by Purkinje neurons and that endogenous myosin 18Aα localizes predominantly to Purkinje neuron spines.

### Expressed, tagged myosin 18Aα localizes to and is enriched in Purkinje neuron spines along with F-actin

Efforts to define the function of myosin 18Aα in Purkinje neurons would be facilitated if expressed versions of the myosin localize within dendritic spines like the endogenous protein. To test this, we co-transfected Purkinje neurons with mCherry as a fill marker and myosin 18Aα tagged at its N-terminus with GFP. Figure 2, Panel A1 through A3 show a representative transfected Purkinje neuron while Panels A4 through A6 show enlarged images of the individual channels and the overlaid image for the boxed region in Panel A3. What is immediately apparent is that expressed, GFP-tagged myosin 18Aα also localizes dramatically to spines. Previous work from our lab using F-Tractin, a live-cell, indirect reporter for F-actin, showed that spines are the major actin-rich compartment in Purkinje neurons (Wagner, Brenowitz and Hammer, 2011). Consistently, Purkinje neurons transfected with GFP-tagged F-Tractin and mCherry-tagged myosin 18Aα show clear co-localization of the two signals within spines (Figure 2; Panels B1 through B3 show the individual channels and overlaid image for a portion of a Purkinje neuron dendritic arbor, Panels B4 through B6 show enlarged images of the boxed region in Panel B3, and Panels B7 through B9 show enlarged images of the boxed region in Panel B6). To determine the fold-enrichment of myosin 18Aα and F-Tractin in spines over dendrites, GFP-tagged myosin 18Aα and GFP-tagged F-Tractin were separately co-transfected with mCherry as a volume marker and ratio imaging was performed to determine the actual difference in the concentrations of each protein in spines relative to the dendrite proper by correcting for differences in the volumes imaged (see Methods). Consistent with the images in Figure 2, myosin 18Aα and F-Tractin are concentrated 3.47 ± 1.11 fold and 3.67 ± 0.99 fold in spines relative to dendrites, respectively (see Figure 4; Panels B1 through B3 and C1 through C3 show representative images for the mCherry/GFP-F-Tractin pair and the mCherry/GFP-myosin 18Aα pair, respectively; Panels A1 through A3 show a representative image of the free mCherry/free GFP control pair; Panel H shows the cumulative results for these three samples). We conclude, therefore, that expressed, tagged myosin 18Aα can be used to interrogate the function of this myosin within Purkinje neuron spines, and that both myosin 18Aα and F-actin are concentered ∼3.5 fold in spines relative to dendrites.

**Figure 2.**
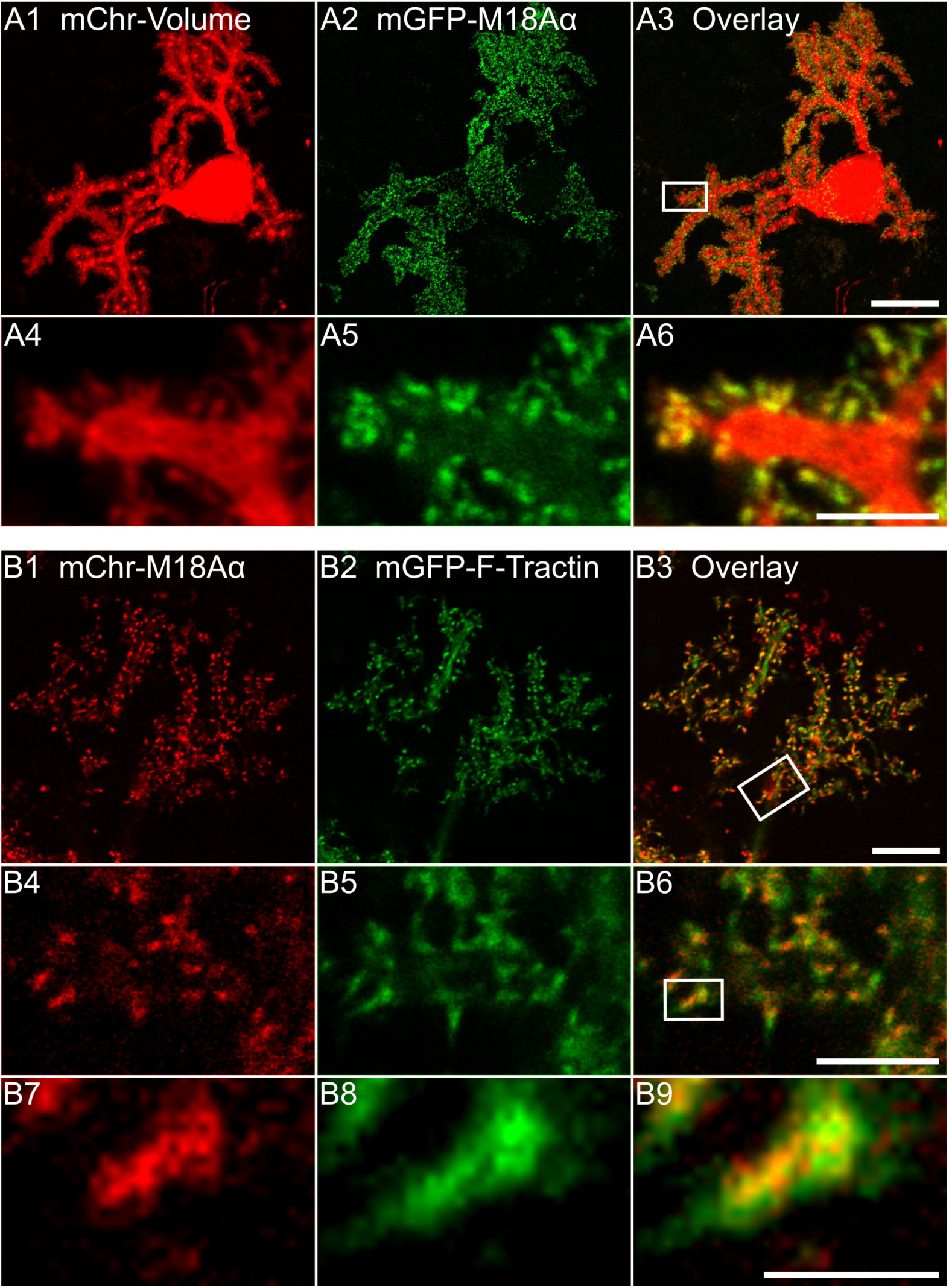
Exogenously expressed myosin 18Aα localizes to dendritic spines together with F-actin. (A1-A6) Panels A1-A3 show a representative cultured Purkinje neuron (DIV 18) that was expressing mCherry as a volume marker (A1) and mGFP-myosin 18α (A2); the overlaid image is shown in Panel A3. Panels A4-A6 show enlargements of the boxed region in Panel A3. (B1-B9) Panels B1-B3 show a portion of the dendritic arbor of a representative cultured Purkinje neuron (DIV 18) that was expressing mCherry-myosin 18Aα (B1) and mGFP-F-Tractin (B2); the overlaid image is shown in (B3). Panels B4-B6 show enlargements of the boxed region in Panel B3. Panels B7-B9 show enlargements of the boxed region in Panel B6. Scale bars: 20 μm (A3), 10 μm (B3), 5 μm (A6 and B6) and 1 μm (B9).

### Myosin 18Aα is enriched towards the base of spines along with myosin 2

To look for evidence that myosin 18Aα might be enriched within a specific region of the spine, we transfected neurons with mCherry-tagged myosin 18Aα and with GFP-tagged PSD93 to mark post synaptic densities. Figure 3, Panels A1 through A3, show the individual channels and overlaid image for a portion of a representative Purkinje neuron dendritic arbor, while Panels A4 through A6 show enlarged images of the boxed region in Panel A3. What is evident, especially from the enlarged images, is that these two signals are largely non-overlapping within spines. Consistently, line scans of fluorescent intensities taken from the base to the tip of spines shows that myosin 18Aα is enriched towards the spine base (Figure 3, Panel D, red curve), while PSD93 is enriched towards the spine tip (Figure 3, Panel D, orange curve). Notably, lines scans of F-Tractin intensities made using images like those in Figure 4, Panels B1 through B3, indicate that F-actin is also enriched towards the spine base, although it usually extends further towards the spine tip than myosin 18Aα (Figure 3, Panel D, blue curve). Together, these light microscopic images argue that myosin 18Aα is enriched towards the base of spines, consistent with the EM images in Figure 1.

**Figure 3.**
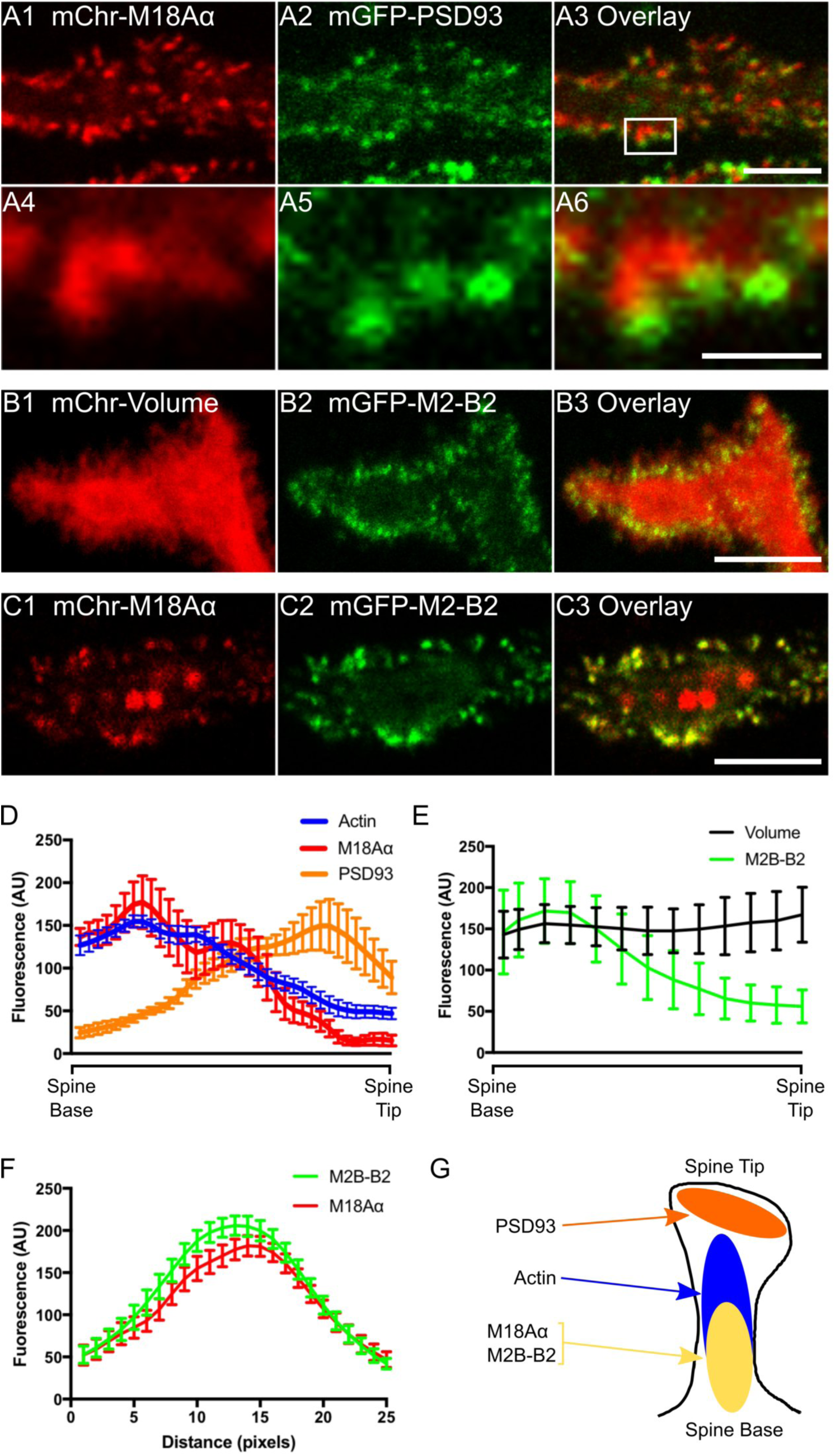
Myosin 18Aα localizes towards the base of the dendritic spine along with myosin 2B. (A1-A6) Panels A1-A3 show a portion of a dendrite from a representative cultured Purkinje neuron (DIV 18) that was expressing mCherry-myosin 18Aα (A1) and mGFP-PSD93 (A2); the overlaid image is shown in (A3). Panels A4-A6 show enlargements of the boxed region in Panel A3. (B1-B3) Shown is a portion of a dendrite from a representative cultured Purkinje neuron (DIV 18) that was expressing mCherry as a volume marker (B1) and mGFP-myosin 2B-B2 (B2); the overlaid image is shown in (B3). (C1-C3) Shown is a portion of a dendrite from a representative cultured Purkinje neuron (DIV 18) that was expressing mCherry-myosin 18Aα (C1) and mGFP-myosin 2B-B2 (C2); the overlaid image is shown in (C3). (D) Shown are the fluorescence intensities (in arbitrary units; AU) from spine base to spine tip for F-actin (blue; determined using mGFP-F-Tractin), myosin 18Aα (red; determined using mCherry-myosin 18Aα), and PSD-93 (orange; determined using mGFP-PSD93) (each plot represents data from at least 20 spines). (E) Shown are the fluorescence intensities (in arbitrary units; AU) from spine base to spine tip for volume (black; determined using an mCherry fill marker) and myosin 2B-B2 (green; determined using mGFP-myosin B2-2B) (each plot represents data from at least 20 spines). (F) Shown are the fluorescence intensities for myosin 2B-B2 (green; determined using mGFP-myosin B2-2B) and myosin 18Aα (red; determined using mCherry-myosin 18Aα) from line scans performed along the long axis of spines (each plot represents data from at least 20 spines). (G) Cartoon depicting the distribution within a typical spine of PSD93 (orange), F-actin (blue), myosin 2B-B2 (yellow), and myosin 18Aα (yellow). Scale bars: 5 μm (A3), 1 μm (A6), 5 μm (B3 and C3).

**Figure 4.**
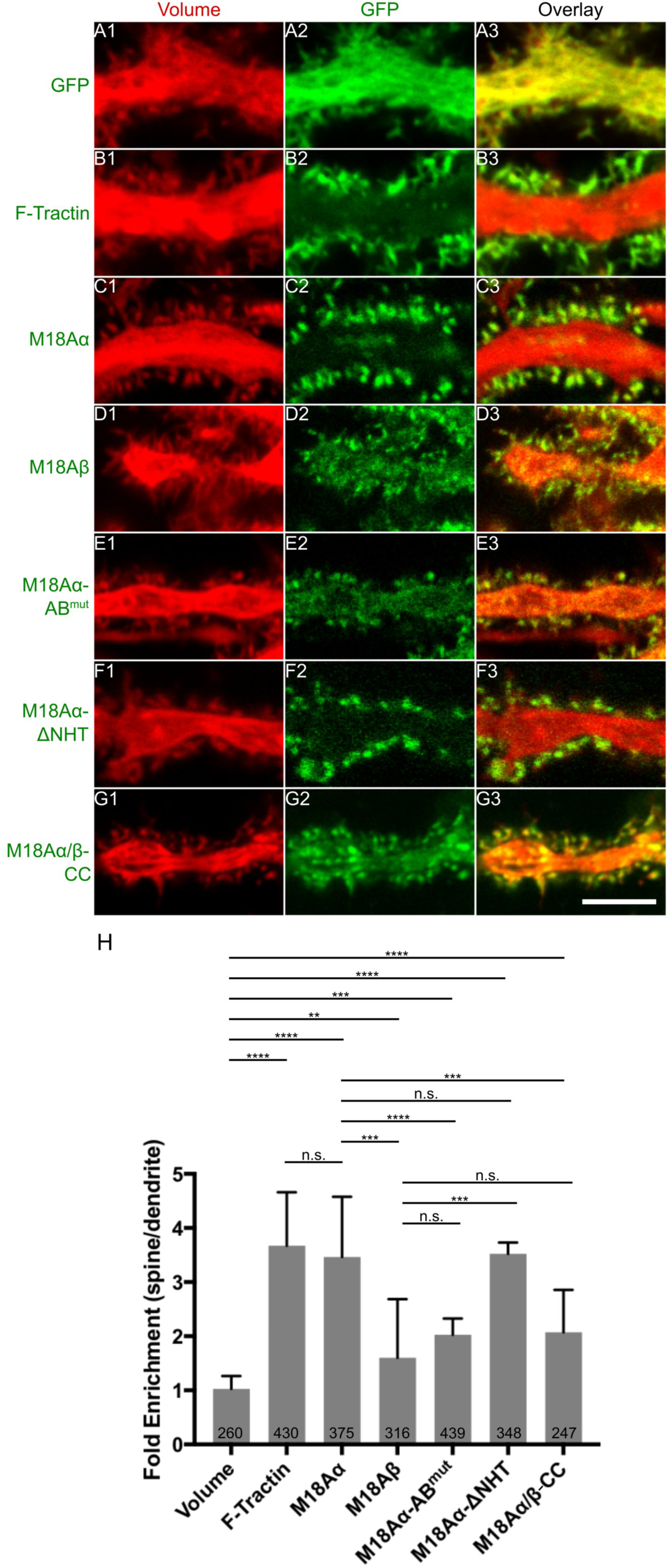
Myosin 18Aα is targeted to spines through a combination of its N-terminal F-actin binding site and co-assembly with myosin 2. Shown are portions of dendrites from representative cultured Purkinje neurons (DIV 18) that were expressing mCherry as a volume marker (A1-G1) and one of the following mGFP-tagged proteins: mGFP alone (A2), F-Tractin (B2), myosin 18Aα (C2), myosin 18Aβ (D2), myosin 18 Aα-AB^mut^ (N-terminal actin binding site mutated) (E2), myosin 18 Aα-ΔNHT (missing non-helical tailpiece) (F2), or myosin 18 Aα/β-CC (coiled coil domain only) (G2). Panels A3-G3 show the overlaid images. (H) Shown are the fold-enrichments in the spine over the adjacent dendrite for each GFP construct, as determined by ratiometric imaging. The number of spines analyzed for each construct, which were derived from three independent experiments, is indicated within each bar. Scale bar: 5 μm (G3). ** *p*<0.01; *** *p*<0.001; **** *p*<0.0001; n.s., not significant.

Previous studies have shown that myosin 2B is enriched towards the base of Hippocampal neuron spines (Morales and Fifková, 1989; Korobova and Svitkina, 2010; Koskinen *et al*., 2014). A similar localization for myosin 2B in Purkinje neuron spines would be consistent with the general thinking that actomyosin contractile activity localizes towards the base of spines, and more specifically with the fact that myosin 18Aα co-assembles with myosin 2 (Billington *et al*., 2015). To address this, we co-transfected Purkinje neurons with mCherry as a volume marker and the B2-spliced isoform of myosin 2B tagged at its N-terminus with GFP (this myosin 2B isoform is Purkinje neuron-specific; (Ma *et al*., 2006)). Figure 3, Panels B1 through B3, show the individual channels and overlaid image for a portion of a representative Purkinje neuron dendritic arbor. As with myosin 18Aα and F-Tractin, myosin 2B-B2 is clearly enriched in spines. Consistently, ratio imaging indicated that it is concentrated 2.7 ± 1.2 fold in spines over dendrites (Figure 4, Panel H). Moreover, lines scans show that myosin 2B-B2 is enriched towards the base of spines like myosin 18Aα (Figure 3, Panel E, green line). Finally, Purkinje neurons expressing both mCherry-tagged myosin 18Aα and GFP-tagged myosin 2B-B2 show strong co-localization of the two myosins within spines (Figure 3, Panels C1 through C3) that is borne out using line scans (Figure 3, Panel F). We conclude, therefore, that myosin 2B-B2 and myosin 18Aα colocalize towards the base of Purkinje neuron spines where they likely co-assemble to form mixed bipolar filaments (Figure 3, Panel G).

### Myosin 18Aα is targeted to spines by a combination the F-actin binding domain present in its N-terminal extension and its ability to co-assemble with myosin 2

To define the motifs within myosin 18Aα responsible for its targeting to spines, we first looked at the targeting of the myosin 18Aβ isoform to spines as a proxy for the role of myosin 18Aα’s N-terminal extension because myosin 18Aβ differs from myosin 18Aα only in lacking this extension (Figure 1A). For this and all subsequent efforts, we performed ratio imaging using a volume marker to obtain values for the concentration of the protein in the spine relative to the dendrite that was corrected for differences in the volumes imaged. Panels D1-D3 and Panel H in Figure 4 together show that while myosin 18Aβ is concentrated to a significant extent in spines, its targeting is less robust than that of myosin 18Aα (1.60 ± 1.10 fold versus 3.47 ± 1.11 fold for myosin 18Aα). This result argues that myosin 18Aα’s N-terminal extension is contributing significantly to spine targeting. To gain further support for this conclusion, we measured the targeting of myosin 18Aα’s isolated N-terminal extension, referred to here as NT. Panels A1 through A3 and Panel E in Figure S3 together show that NT is indeed enriched in spines 3.26 ± 1.44 fold over dendrites.

The NT sequence contains three recognizable domains: a KE-rich region, a nucleotide-independent F-actin binding site (Taft *et al*., 2013), and a PDZ domain (Tonikian *et al*., 2008) (Figure 1A). To define the relative importance of these three domains for spine targeting, we measured the targeting of NT constructs in which (i) the N-terminal, 29-residue KE-rich region was deleted (NT-ΔKE), (ii) a function blocking point mutation (Tonikian *et al*., 2008) was introduced into the PDZ domain (NT-PDZ^mut^), or (iii) two point mutations were introduced into the F-actin binding site that together reduce its affinity for F-actin *in vitro* by ∼10-fold (NT-AB^mut^; see Figure S4 for the F-actin binding assays that were performed using NT and NT-AB^mut^). Panels B1 through B3, C1 through C3, D1 through D3 and Panel E in Figure S3 together show that only the mutation of the F-actin binding site in NT significantly reduced its spine targeting (from 3.26 ± 1.44 fold for NT to 1.71 ± 0.81 fold for NT-AB^mut^). Consistent with this observation, Panels E1 through E3 and Panel H in Figure 4 together show that full length myosin 18Aα containing this F-actin binding site mutation exhibited a significant reduction in spine targeting (2.03 ± 0.35 fold versus 3.47 ± 1.11 fold for wild type myosin 18Aα).

While the preceding results clearly indicate that the nucleotide-independent F-actin binding site present within M18Aα’s N-terminal extension contributes to its spine targeting, the spine targeting that persists when this binding site is impaired or absent indicates that another motif(s) within myosin 18Aα must be contributing significantly to targeting. The two most obvious remaining candidates are the PDZ ligand comprising the C-terminal 7 residues of myosin 18Aα, and myosin 18Aα’s coiled-coil domain, which drives its co-assembly with myosin 2 (Billington *et al*., 2015). Panels F1 through F3 and Panel H in Figure 4 together show that deletion of myosin 18Aα’s 90-residue non-helical tailpiece (ΔNHT), which contains the PDZ ligand, does not have any adverse effect on the targeting of myosin 18Aα to spines. In contrast, Panels G1 though G3 and Panel H in Figure 4 show that myosin 18Aα’s coiled-coil domain exhibits significant spine targeting (2.07 ± 0.78 fold spine enrichment versus 3.47 ± 1.11 fold for full length myosin 18Aα). Based on all of these results, we conclude that the targeting of myosin 18Aα to Purkinje neuron spines is driven by a combination of the F-actin binding site present within its N-terminal extension and its ability to co-assemble with myosin 2 via its coiled-coil domain.

### Myosin 18A knockdown results in increases in spine length and density consistent with an impairment in spine maturation

Having established that myosin 18Aα targets robustly to Purkinje neuron spines, we next asked what effect the loss of this myosin might have on spine development. To address this question, we made use of a Purkinje neuron-specific miRNA-mediated knockdown system we recently developed (Alexander and Hammer, 2016). Like the custom plasmids we use to express genes specifically within Purkinje neurons (Wagner, Brenowitz and Hammer, 2011), this knockdown system makes use of the Purkinje neuron-specific promoter L7/Pcp2 to drive miRNA expression. Validation of this system was obtained previously (Alexander and Hammer, 2016) by showing that wild type Purkinje neurons expressing a miRNA directed against myosin Va exhibit the same phenotype as Purkinje neurons isolated from mice homozygous for a myosin Va functional null allele (loss of inheritance of endoplasmic reticulum into spines) (Wagner, Brenowitz and Hammer, 2011). Because the efficiency of knockdown within cultured Purkinje neurons cannot be accessed by Western blotting (as they represent a minor fraction of the cells present in mixed primary cultures), we first measured the extent of myosin 18Aα knockdown in NIH/3T3 fibroblasts using two candidate miRNAs directed against the 3’ untranslated region of myosin 18Aα. Both miRNAs yielded ∼80% knockdown of myosin 18Aα (Figure S5, lanes 1-3). Given this, we introduced miRNA #2 into wild type Purkinje neurons at day 0 using our Purkinje neuron-specific knockdown plasmid. Of note, this plasmid also drives the expression of free mCherry, which serves to identify Purkinje neurons expressing the miRNA and to mark the cell’s volume. As a control, we used a scrambled miRNA (Alexander and Hammer, 2016). Consistent with penetrant knockdown, antibody staining of wild type Purkinje neurons expressing the scrambled miRNA (Figure S5, Panels A1-A3) side by side with Purkinje neurons expressing miRNA #2 (Figure S5, Panels B1-B3) after 18 days in culture revealed a large decease in the signal for myosin 18Aα within the spines of the miRNA #2-treated neurons.

Live-cell imaging of control Purkinje neurons expressing the scrambled miRNA (Figure 5, Panels A1 through A4) and Purkinje neurons expressing myosin 18Aα miRNA #2 (Figure 5, Panels B1 and B2) after 18 days in culture revealed an apparent difference between them in terms of spine length, with the knockdown neurons appearing to exhibit longer spines (see also the higher magnification images in Panel E (scrambled miRNA) and Panel F (M18Aα miRNA)). Indeed, quantitation (Figure 5, Panel I) showed that myosin 18Aα knockdown Purkinje neurons exhibit about a 30% increase in spine length relative to control neurons (1.32 ± 0.39 µm for knockdown neurons versus 1.03 ± 0.29 µm for control neurons; note that this later value matches values reported previously (Heintz, Eva and Fawcett, 2016) for the length of spines on mature wild type Purkinje neurons present in primary culture). More generally, the spines on the myosin 18Aα knockdown Purkinje neurons appear more filopodial in nature (Figure5, Panels E and F), as if the maturation of filopodial precursors into mature spines (Yuste and Bonhoeffer, 2004; Ethell and Pasquale, 2005; Tada and Sheng, 2006; Lee *et al*., 2007; Yoshihara, De Roo and Muller, 2009; Gao *et al*., 2011; Velázquez-Zamora, Martínez-Degollado and González-Burgos, 2011; Kanjhan, Noakes and Bellingham, 2016) is impaired by the knockdown of myosin 18Aα. Quantitation (Figure 5, Panel J) also showed that myosin 18Aα knockdown Purkinje neurons exhibit about a 1.7-fold increase in the density of spines relative to control neurons (3.43 ± 1.33 spines per µm^2^ for knockdown neurons versus 2.05 ± 1.1 spines per µm^2^ for control neurons). This phenotype may also reflect an impairment in spine maturation, given that the spines of myosin 18Aα knockdown Purkinje neurons contain less myosin 2B ((Hodges *et al*., 2011)es; see also below and Discussion). Importantly, these defects in spine length and density were both completely rescued by reintroduction of wild type, miRNA-resistant myosin 18Aα into myosin 18Aα knockdown Purkinje neurons (see Figure 6, Panels C1 though C4, for a representative rescued Purkinje neuron, Panel G for a higher magnification image of a dendrite from one such neuron, and Panels I and J for quantitation of spine length and density; spine length: 1.00 ± 0.28 µm; spine density: 2.05 ± 1.10 spines per µm^2^). This result argues that the apparent defect in spine maturation exhibited by Purkinje neurons expressing the myosin 18Aα miRNA is due to the knockdown of myosin 18Aα and not to possible off-target effects. We also note that 68.2 ± 8.8% of spines on myosin 18Aα knockdown Purkinje neurons had a closely associated Bassoon-positive puncta, which is not significantly different from the value for control neurons (73.7 ± 3.3%). This observation argues that the defect in spine maturation exhibited by myosin 18Aα knockdown Purkinje neurons is not caused by a reduction in the fraction of spines that are active.

**Figure 5.**
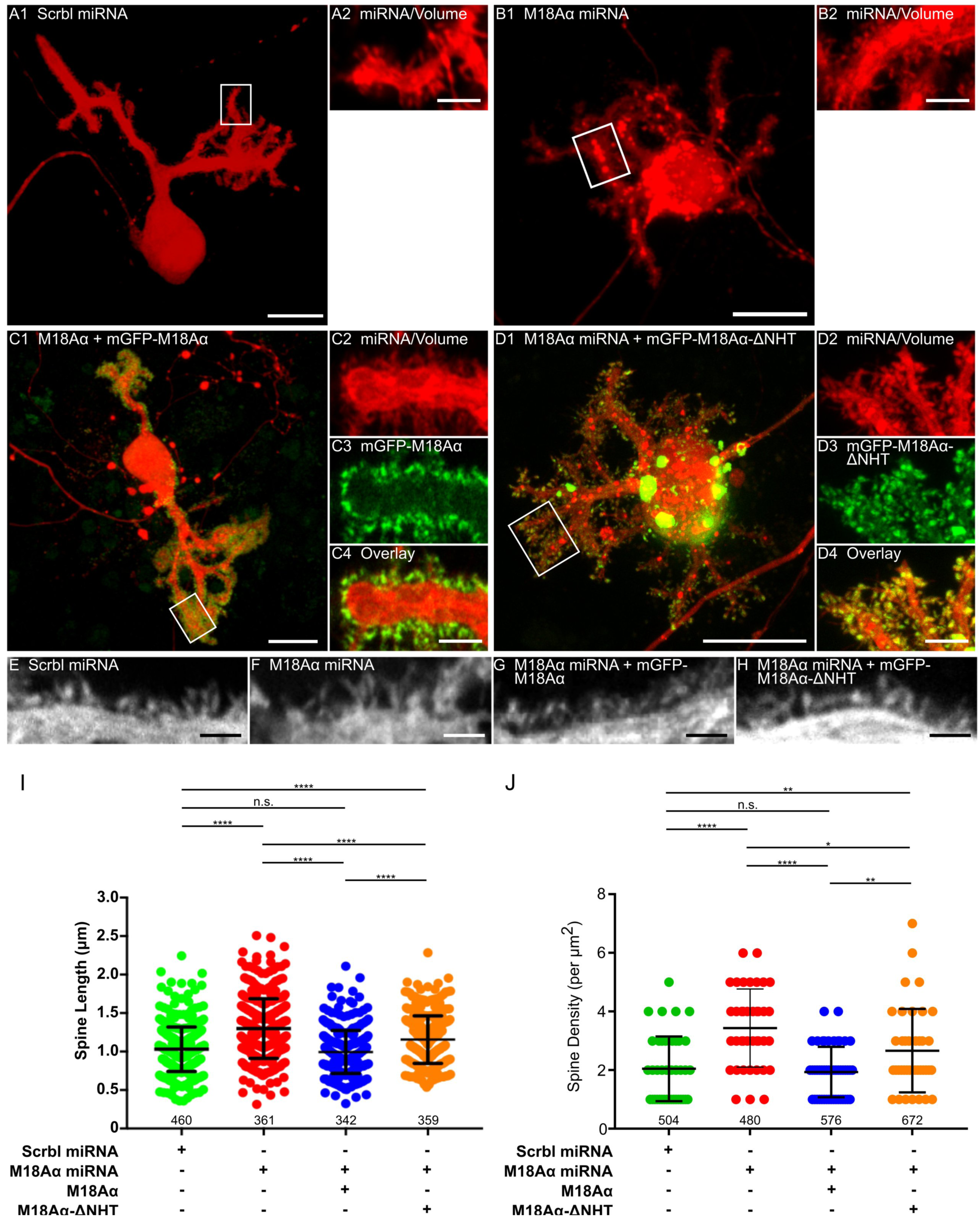
Knockdown of myosin 18Aα in Purkinje neurons results in an increase in spine length and density, consistent with a defect in spine maturation. (A1 and A2) Panel A1 shows a representative cultured Purkinje neuron (DIV 18) that was expressing a scrambled miRNA (indicated by the presence of the mCherry volume marker). Panel A2 shows an enlargement of the boxed region in Panel A1. (B1 and B2) As in A1 and A2, except the Purkinje neuron was expressing myosin 18 Aα miRNA #2. (C1-C4) Panel C1 shows a representative cultured Purkinje neuron (DIV 18) that was expressing myosin 18 Aα miRNA #2 (indicated by the presence of the mCherry volume marker) and mGFP-myosin 18 Aα. Panels C2-C4 show enlargements of the boxed region in Panel C1. (D1-D4) Panel D1 shows a representative cultured Purkinje neuron (DIV 18) that was expressing myosin 18 Aα miRNA #2 (indicated by the presence of the mCherry volume marker) and mGFP-myosin 18 Aα-ΔNHT. Panels D2-D4 show enlargements of the boxed region in Panel D1. Panels E-H show representative, high-magnification images of spines on Purkinje neurons treated as in Panels A1, B1, C1, and D1, respectively. (I) Measurements of spine length (in µm) on DIV 18 Purkinje neurons treated as indicated (green, scrambled miRNA; red, myosin 18Aα miRNA; blue, myosin 18Aα miRNA plus GFP-myosin 18Aα; orange, myosin 18Aα miRNA plus GFP-myosin 18Aα-ΔNHT). The N values, which correspond to the total number of spines scored using at least 10 neurons across three independent experiments, are indicated below each measurement. (J) Measurements of spine density (in numbers per µm^2^) on DIV 18 Purkinje neurons treated as indicated. The N values, which correspond to the total area scored using at least 10 neurons from three independent experiments, are indicated below each measurement. Scale bars: 20 μm (A1, B1, C1 and D1), 5 μm (A2, B2, C4 and D4), 1 μm (E-H). * *p*<0.05; ** *p*<0.01; **** *p*<0.0001; n.s., not significant.

**Figure 6.**
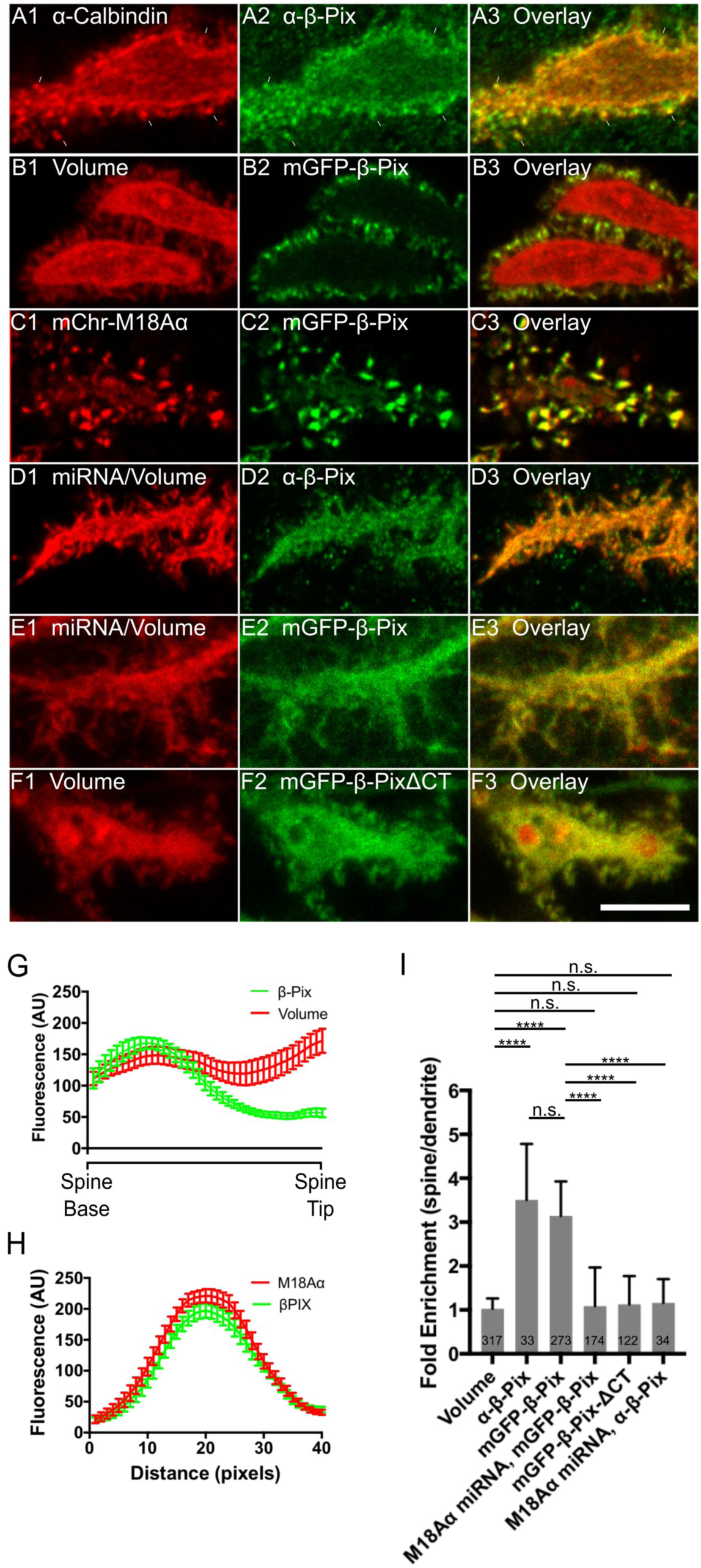
Myosin 18Aα targets β-Pix to Purkinje neuron spines. Shown are portions of dendrites from representative cultured Purkinje neurons (DIV 18) treated as follows. (A1-A3) Fixed and stained for Calbindin (A1) and endogenous β-Pix (A2); Overlay (A3). (B1-B3) Expressing mCherry fill marker (B1) and mGFP-β-Pix (B2); Overlay (B3). (C1-C3) Expressing mCherry-myosin 18Aα (C1) and mGFP-β-Pix (C2); Overlay (C3). (D1-D3) Expressing myosin 18 Aα miRNA (D1; associated mCherry fill marker shown) and fixed/stained for endogenous β-Pix (D2); Overlay (D3). (E1-E3) Expressing myosin 18 Aα miRNA #2 (E1; associated mCherry fill marker shown) and mGFP-β-Pix (E2); Overlay (E3). (F1-F3) Expressing mCherry fill marker (F1) and mGFP-β-PixΔCT (F2); Overlay (F3). (G) Shown are the fluorescence intensities (in arbitrary units; AU) from spine base to spine tip for volume (red; determined using an mCherry fill marker) and β-Pix (green; determined using mGFP-β-Pix) (each plot represents data from at least 20 spines). (H) Shown are the fluorescence intensities for myosin 18Aα (red; determined using mCherry-myosin 18Aα) and β-Pix (green; determined using mGFP-β-Pix) from line scans performed along the long axis of spines (each plot represents data from at least 20 spines). (I) Shown are the fold-enrichments in the spine over the adjacent dendrite for each GFP construct or stained protein, as determined by ratio imaging. The number of spines analyzed for each construct or stained protein is indicated within each bar. A total of 6 cells from three independent experiments were analyzed for each GFP construct. Scale bar: 5 μm (F3). **** *p*<0.0001; n.s. not significant.

To gain further support for the results obtained using miRNA-mediated myosin 18Aα knockdown, we examined Purkinje neurons isolated from a myosin 18A conditional knockout (cKO) mouse following the delivery of Cre recombinase into these neurons using TAT-Cre. While the creation and characterization of this mouse will be described elsewhere, we show in Figure S5 (Panel D) that MEFs isolated from this mouse and transfected with a plasmid expressing Cre recombinase lose essentially all of the signal in a Western blot for both myosin 18Aα and myosin 18Aβ, indicating that the conditional allele we created is functional. Figure S6 shows representative images at day 18 in culture (with enlarged insets) of a cKO Purkinje neuron that was not treated with TAT-Cre (Panel A), a cKO Purkinje neuron that was treated TAT-Cre at day 5 (Panel B), and a wild type Purkinje neuron that was treated with TAT-Cre at day 5 (Panel C). Quantitation (Figure S6, Panels D and E) shows that cKO Purkinje neurons treated with TAT-Cre exhibit significant increases in both spine length (0.92 ± 0.22 µm for cKO neurons without TAT-Cre treatment versus 1.28 ± 0.35 µm for cKO Purkinje neurons with TAT-Cre treatment) and spine density (2.95 ± 1.38 spines per µm^2^ for cKO neurons without TAT-Cre treatment versus 4.08 ± 2.14 spines per µm^2^ for cKO Purkinje neurons with TAT-Cre treatment). Importantly, the length and density of spines on wild type Purkinje neurons treated at day 5 with TAT-Cre (Figure S6, Panel C) was not significantly different from cKO Purkinje neurons not treated with TAT-Cre (Figure S6, Panels D and E; spine length: 0.93 ± 0.28 µm; spine density: 2.51 ± 1.24 spines per µm^2^), indicating that the effect of TAT-Cre on spine length and density in cKO Purkinje neurons is likely due to the loss of myosin 18Aα and not to some non-specific effect caused by the introduction of Cre recombinase. Together, these results, like the knockdown results described above, argue that myosin 18Aα plays a significant role in the maturation of Purkinje neuron spines.

An additional phenotype arising from the knockdown/knockout of myosin 18Aα that is seen in all of the related images discussed above (compare, for example, Panel B1 to Panel A1 in Figure 5) is an apparent defect in cell polarity manifested as an increase in the number of primary dendrites in myosin 18Aα knockdown/knockout neurons. Consistently, Figure S7 shows that while wild type Purkinje neurons, control Purkinje neurons expressing the scrambled miRNA, cKO Purkinje neurons not treated with TAT-Cre, and wild type Purkinje neurons treated with TAT-Cre all exhibit about two primary dendrites per neuron (2.0 ± 0.9, 1.9 ± 0.7, 2.50 ± 1.0, and 1.7 ± 0.8, respectively), myosin 18Aα knockdown Purkinje neurons and cKO Purkinje neurons treated with TAT-Cre exhibit significantly higher numbers of primary dendrites per neuron (5.6 ± 2.2 and 4.00 ± 1.7 primary dendrites per neuron, respectively). Importantly, myosin 18Aα knockdown Purkinje neurons in which wild type myosin 18Aα had been reintroduced exhibit normal numbers of dendrites per cell (2.1 ± 0.8 primary dendrites per neuron; Figure S7), arguing that the polarity defect is due to the loss of myosin 18Aα. While we did not pursue the underlying mechanism of this defect in dendrite morphogenesis, it may be related to the reported interaction of myosin18A with Golgi membranes ((Dippold *et al*., 2009); also see Discussion).

### Myosin 18Aα targets the Rac/Cdc42 GEF β-Pix to Purkinje neuron spines

We next sought to gain insight into the mechanism by which myosin 18Aα contributes to spine maturation. As discussed in the Introduction, myosin 18A isoforms probably contribute to cell function primarily by co-assembling with myosin 2, where they then recruit proteins that regulate the myosin filament, influence the local environment around the filament, and/or attach the filament to cellular structures. One protein known to interact with myosin 18Aα is the Rac/Cdc42 GEF β-Pix, which binds to the myosin’s C-terminal non-helical (Hsu *et al*., 2010, 2014). This interaction could be particularly relevant here, as β-Pix has been implicated in promoting spine maturation via several distinct pathways involving actin assembly and myosin assembly and contractility (see Introduction). Given this, we asked if myosin 18Aα’s contribution to spine maturation is mediated at least in part by its interaction with β-Pix.

We first asked if β-Pix is targeted to Purkinje neuron spines. Figure 6, Panels A1 through A3, show that endogenous β-Pix appears enriched in spines (see arrowheads). Consistently, ratio imaging indicated that it is concentrated 3.51 ± 1.27 fold in spines over dendrites (Figure 6, Panel I). Figure 6, Panels B1 through B3, show that GFP-tagged β-Pix is also enriched is spines, with ratio imaging indicating that it is concentrated 3.14 ± 0.79 fold in spines over dendrites (Figure 6, Panel I). Moreover, line scans of individual spines showed that, like myosin 18Aα and myosin 2, β-Pix is enriched towards the base of the spine (Figure 6, Panel G). Consistently, mCherry-tagged myosin 18Aα and mGFP-tagged β-Pix colocalize dramatically in spines (Figure 6, Panels C1 through C3, and Panel H). Together, these results show that β-Pix is targeted to Purkinje neuron spines and colocalizes with myosin 18Aα.

We next asked if the targeting of β-Pix to spines depends on myosin 18Aα. Figure 6, Panels D1 through D3, show that the enrichment of endogenous β-Pix in spines appears to be lost upon myosin 18Aα knockdown (and, as predicted, the spines appear more filopodia-like). Consistently, ratio imaging indicated that endogenous β-Pix is indeed no longer concentrated in spines relative to dendrites following myosin 18Aα knockdown (Figure 6, Panel I). Similar results were obtained using GFP-tagged β-Pix, which is also no longer concentrated in spines relative to dendrites following myosin 18Aα knockdown (Figure 6, Panels E1 through E3, and Panel I). Finally, a GFP-tagged version of β-Pix lacking C-terminal residues 639-647 (β-Pix-ΔCT), which is no longer able to bind to myosin 18Aα (Hsu *et al*., 2010, 2014), does not exhibit significant spine enrichment (Figure 6, Panels F1 through F3, and Panel I). Together, these results argue that the recruitment of β-Pix to Purkinje neuron spines depends on myosin 18Aα.

### The spines of myosin 18Aα knockdown neurons exhibit significant reductions in the content of both F-actin and myosin 2

The above results are consistent with the idea that the defect in spine maturation exhibited by Purkinje neurons lacking myosin 18Aα might be caused, at least in part, by the loss of β-Pix targeting to spines. Based on previous studies (see Introduction), β-Pix can promote spine maturation downstream of its GEF activity towards Rac1 and Cdc42 via at least two pathways: increased Arp2/3 complex-dependent branched actin filament nucleation and increased myosin 2 filament assembly/contractility following the PAK-dependent phosphorylation of the myosin’s RLC. In our particular case, this latter pathway could be driven by the myosin 18Aα-dependent recruitment of PAK to spines (as it is commonly in a complex with β-Pix and GIT1; see Discussion), as well as by the activation of PAK downstream of β-Pix’s GEF activity. To look for evidence that the defect in spine maturation exhibited by Purkinje neurons lacking myosin 18Aα is due at least in part to defects in these two pathways, we measured the spine content of F-actin and myosin 2 in control and myosin 18Aα knockdown neurons by staining cells with either Phalloidin or an antibody to myosin 2B, respectively. Figure 7, Panels A1-A3, show a representative example of a dendrite from a control Purkinje neuron (DIV 18) that was fixed and stained for Calbindin and F-actin, while Panels B1-B3 show a representative example of a dendrite from a Purkinje neuron that was treated with myosin 18Aα miRNA #2 (DIV 18) and then fixed and stained for F-actin. Comparison of the images in Panels A2 and B2 suggested that myosin 18Aα knockdown results in a significant reduction in Phalloidin staining/F-actin content per spine. Consistently, quantitation showed that knockdown neurons exhibit on average a ∼35% reduction in total F-actin content per spine relative to control neurons (Figure 7, Panel C).

**Figure 7.**
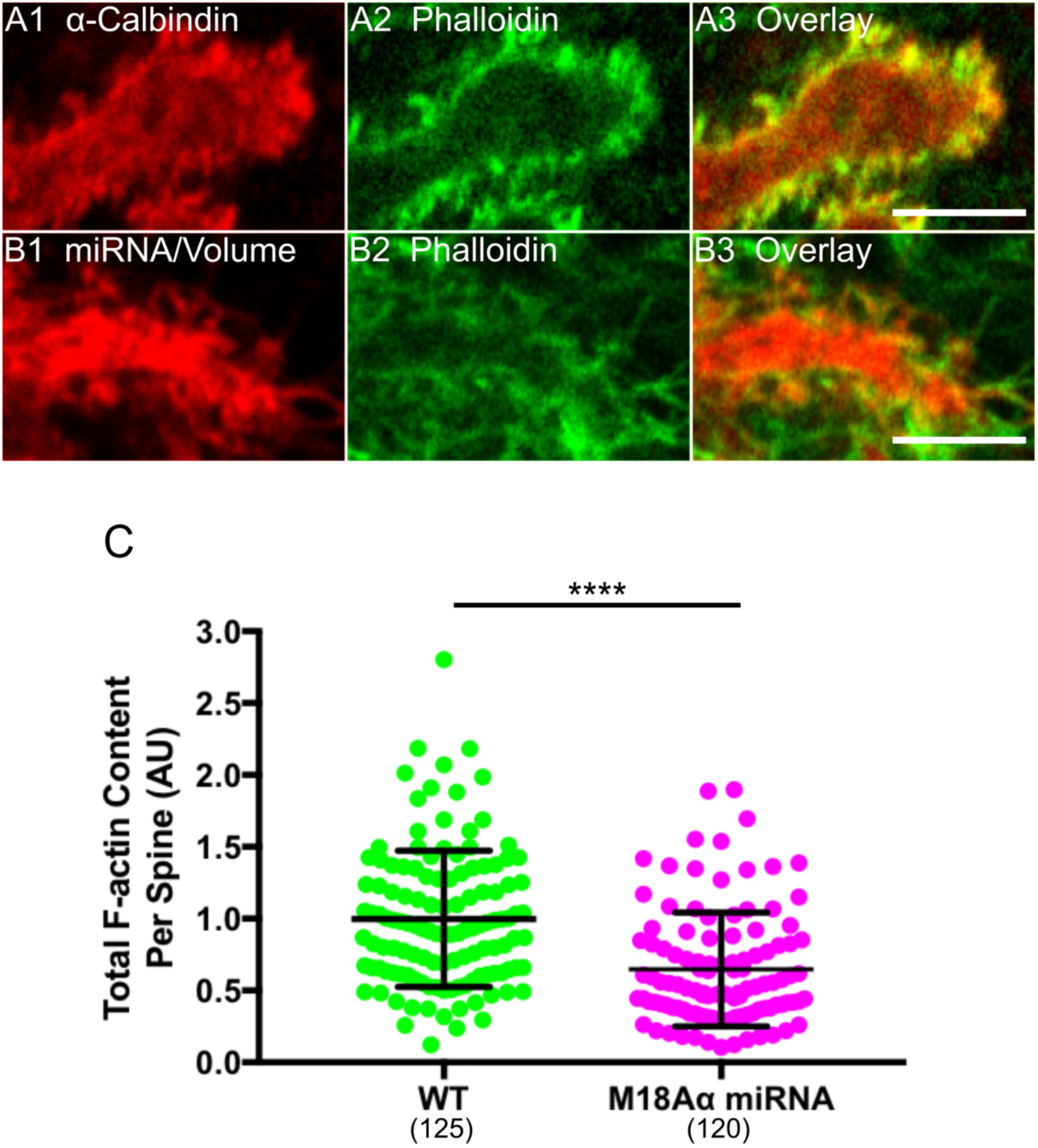
Myosin 18Aα knockdown neurons exhibit reduced F-actin content per spine. (A1-A3) Shown is a portion of a dendrite from a representative cultured control Purkinje neuron (DIV 18) that was fixed and stained for Calbindin (A1) and F-actin using Phalloidin (A2); Overlay (A3). (B1-B3) Shown is a portion of a dendrite from representative cultured Purkinje neuron (DIV 18) that was expressing myosin 18 Aα miRNA #2 (indicated by the presence of the mCherry volume marker) (B1) and then fixed and stained for F-actin using Phalloidin (B2); Overlay (B3). (C) Phalloidin fluorescence/F-actin content per spine in arbitrary units (AU) (WT, green, n = 125 spines measured; myosin 18Aα miRNA, magenta, n = 120 spines measured; the results are from two independent experiments). The fluorescence values were normalized such that the mean intensity in WT spines equals 1.0. Of note, the difference in spine F-actin content between WT and knockdown neurons was maintained when the two samples were normalized for spine volume by ratio imaging (34.6% reduction in F-actin content per unit spine volume versus 35.2% reduction in total actin content per spine). Scale bar: 5 μm (A3 and B3). **** *p*<0.0001.

With regard to the content of myosin 2B in spines, Figure 8, Panels A1-A3 (and the corresponding insets in Panels A4-A6), show a representative example of a dendrite from a control Purkinje neuron (DIV 18) that treated with a scrambled miRNA and then fixed and stained for myosin 2B, while Panels B1-B6 show the same except that the Purkinje neuron was treated with myosin 18Aα miRNA #2. Comparison of the images in Panels A2/A5 with those in Panels B2/B5 suggested that myosin 18Aα knockdown results in a significant reduction in myosin 2B content per spine. Consistently, quantitation showed that knockdown neurons exhibit on average a ∼39% reduction in total myosin 2B content per spine relative to control neurons (Figure 8, Panel C). These results, together with those in Figure 7, argue that the defect in spine maturation caused by the loss of myosin 18Aα may be due, at least in part, to an attenuation of the two β-Pix-driven maturation pathways discussed above.

**Figure 8.**
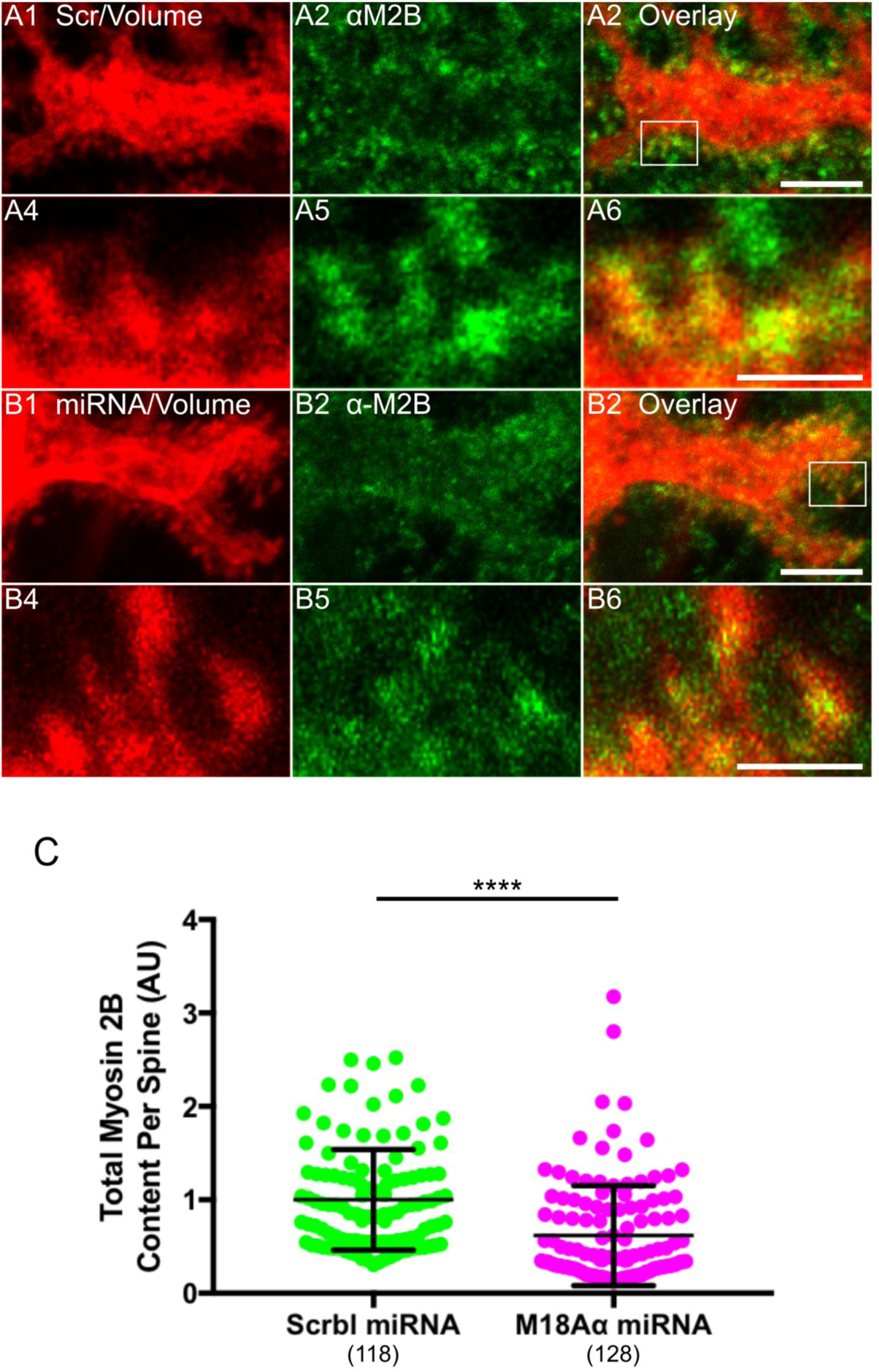
Myosin 18Aα knockdown neurons exhibit reduced myosin 2B content per spine. (A1-A6) Panels A1-A3 show a portion of a dendrite from representative cultured control Purkinje neuron (DIV 18) that was expressing a scrambled miRNA (indicated by the presence of the mCherry volume marker) (A1) and then fixed and stained for myosin 2B (A2); Overlay (A3). Panels A4-A6 show enlargements of the boxed region in Panel A3. (B1-B6) Panels B1-B3 show a portion of a dendrite from representative cultured Purkinje neuron (DIV 18) that was expressing myosin 18 Aα miRNA #2 (indicated by the presence of the mCherry volume marker) (B1) and then fixed and stained for myosin 2B (B2); Overlay (B3). Panels B4-B6 show enlargements of the boxed region in Panel B3. (C) Myosin 2B fluorescence/content per spine in arbitrary units (AU) (Control, green, n = 118 spines measured; myosin 18Aα knockdown, magenta, n = 128 spines measured; the results are from three independent experiments). The fluorescence values were normalized such that the mean intensity in control spines equals 1.0. Of note, the difference in spine myosin 2B content between control and knockdown neurons was maintained when the two samples were normalized for spine volume by ratio imaging (34.6% reduction in myosin 2B content per unit spine volume versus 39.3% reduction in total myosin 2B content per spine). Scale bar: 5 μm (A3 and B3) and 1 μm (A6 and B6). **** *p*<0.0001.

### Myosin 18Aα’s contribution to spine maturation depends significantly on its interaction with β-Pix but extends beyond it as well

If myosin 18Aα’s contribution to spine maturation is due entirely to its ability to recruit β-Pix to spines, then a version of myosin 18Aα lacking its C-terminal non-helical tailpiece (M18Aα ΔNHT), which can no longer bind to β-Pix (Hsu *et al*., 2010, 2014), should not rescue knockdown cells. Somewhat surprisingly, Figure 5, Panels D1 through D4, together with Panels I and J, show that this construct results in a significant degree of rescue of both spine length and spine density. That said, the values for spine length and density using this construct are also significantly different from the values for control cells. In other words, a version of myosin 18Aα that cannot target β-Pix to spines yields a partial rescue of knockdown cells in which the values for spine length and density are in between, and significantly different from, the values for both control and knockdown cells. The simplest interpretation of this result is that while the interaction of myosin 18Aα with β-Pix plays a significant role in driving spine maturation, there must be at least one other pathway by which myosin 18Aα contributes significantly to spine maturation (see Discussion).

## DISCUSSION

Here we show that myosin 18Aα recruits the Rac1/Cdc42 GEF β-Pix to the spines of cerebellar Purkinje neurons to promote spine maturation. With regard to the underlying mechanism of this effect, our demonstration that the spines of myosin 18Aα knockdown Purkinje neurons contain significantly less F-actin and myosin 2B argues that the myosin 18Aα-dependent recruitment of β-Pix serves to enhance the assembly of actin and myosin filaments in spines. This conclusion is consistent with previous studies showing that β-Pix promotes spine maturation in hippocampal neurons downstream of its Rac1/Cdc42 GEF activity by increasing Arp2/3 complex-dependent branched actin filament nucleation (via activation of the NPFs WAVE and WASp) (Kim *et al*., 2006; Soderling *et al*., 2007; Rácz and Weinberg, 2008; Wegner *et al*., 2008; Chen, Wirth and Ponimaskin, 2012; Tejada-Simon, 2015; Spence *et al*., 2016), and by increasing myosin 2 filament assembly/contractility (via the activation of PAK-dependent myosin RLC phosphorylation) (Zhang *et al*., 2003, 2005). Together, our results add additional support to the emerging view that myosin 18A isoforms confer via their N- and C-terminal protein interaction domains novel functions upon myosin 2 by co-assembling with it (Tan *et al*., 2008; Hsu *et al*., 2010, 2014; Billington *et al*., 2015).

One result that surprised us was that myosin 18Aα lacking its non-helical tailpiece where β-Pix binds was able to partially rescue knock down cells with regard to spine length and spine density. This result argues that the interaction of myosin 18Aα with β-Pix, while important, is not the whole story, and that there must be at least one other pathway by which myosin 18Aα contributes significantly to spine maturation. While we did not attempt to define this pathway, a number of candidates come to mind that could guide future work. First, myosin 18Aα could promote spine maturation by stabilizing myosin 2 filaments (Billington *et al*., 2015). Second, myosin 18Aα could promote spine maturation by recruiting the myosin 2 regulatory light chain kinase MRCK via its N-terminal PDZ domain (and through the adaptor protein LRAP; (Tan *et al*., 2008)). Third, other yet-to-be identified proteins that interact with myosin 18Aα’s N-terminal extension could promote spine maturation by promoting actin and/or myosin assembly. Future work will address these and other possible mechanisms by which myosin 18Aα promotes spine maturation beyond its role in recruiting β-Pix to spines.

The only morphological indicator of a defect in spine maturation that we provided at the individual spine level was increased spine length. This is in contrast to studies in cultured hippocampal neurons, where reduced spine head size is usually included along with increased spine length as an indicator of a defect in spine maturation (Harris and Stevens, 1989; Harris, Jensen and Tsao, 1992; Fiala *et al*., 1998; Takahashi *et al*., 2003; Oray, Majewska and Sur, 2005). Unlike hippocampal neurons *in situ*, where ∼70% of mature spines exhibit an enlarged spine head, only ∼10% of mature spines on Purkinje neurons *in situ* exhibit an enlarged spine head (Lee, Kim and Rhyu, 2005). Consistently, we rarely see spines with enlarged heads in cultured Purkinje neurons at DIV18. It is for this reason that our only measure of a defect in spine maturation at the individual spine level was increased spine length.

The other spine parameter we measured was spine density, which increased upon myosin 18Aα knockdown/knockout. This change could reflect an attenuation of spine pruning (Stein and Zito, 2018), as this complex process often parallels spine maturation, although we have no direct evidence that spine pruning slows in knock down neurons. A more likely explanation, however, has to do with our observation that the spines of myosin 18Aα knockdown Purkinje neurons contain significantly less myosin 2B, as attenuation of myosin 2B function in hippocampal neurons leads to an increase in the density of immature, filopodial-like spine precursors (Hodges *et al*., 2011). If this is the case, then the increase in spine density exhibited by myosin 18Aα knockdown Purkinje neurons represents a second morphological indicator of a defect in spine maturation.

We presented two supplementary figures containing data obtained using Purkinje neurons isolated from a myosin 18Aα cKO mouse. The availability of this mouse opens the door to the characterization of myosin 18Aα function at the level of the cerebellum (e.g. its role in synaptic plasticity) and the animal (e.g. its role in coordination and motor learning). That said, when we crossed our myosin 18Aα cKO mouse with the *L7/Pcp2* cre driver mouse, only a small fraction of Purkinje neurons in 3 month-old animals appeared to have lost myosin 18Aα expression based on tissue staining (data not shown). This may be due to issues with the access of Cre to the LoxP sites, although the ability to delete the gene in cKO MEFs (Figure S5) would appear to argue against this. A second issue may be the relative weakness of the *L7/Pcp2* promoter (Sugawara *et al*., 2013). Available alternatives are a *L7/Pcp2* Cre driver mouse created using BAC transgene technology (Zhang *et al*., 2004), or stereotaxic injection of an adenovirus harboring Cre driven by an improved version of the *L7/Pcp2* promoter (Nitta *et al*., 2017).

Our β-Pix targeting data indicates that the bulk of the protein is present in the central and lower regions of Purkinje neuron spines. This differs from older reports in hippocampal neurons that β-Pix, as well as other GEFs (and GAPs) for Rho-related GTPases, are targeted to the PSD (Kiraly *et al*., 2010; Chazeau and Giannone, 2016). Indeed, even key effectors of these signaling molecules like the NPFs for the Arp2/3 complex were reported to be localized at the PSD. While our results for β-Pix may differ because we used Purkinje neurons, it is important to note that more recent studies in hippocampal neurons have provided clear evidence that Rho GTPases, as well as their upstream regulators and downstream effectors, localize away from the PSD (Penzes and Cahill, 2012). This is consistent with the fact that many of the pathways driven by these signaling and effector molecules (e.g. actin assembly and myosin contractility) are operating at considerable distances from the PSD (e.g. in the middle and at the base of spines). Relevant to this, a result showing that β-Pix’s GEF activity is required for the maturation of Purkinje neuron spines would add significantly to our story. Repeated efforts to demonstrate this using knockdown/replacement type experiments were unfortunately unsuccessful (no transfected cells were observed on three attempts). Finally, future efforts should seek to define the extent to which the myosin 18Aα-dependent recruitment of β-Pix to spines also determines the spine localization of GIT1 and PAK, which are usually in a complex with β-Pix (Zhang *et al*., 2003, 2005; Za *et al*., 2006; Kiraly *et al*., 2010).

One additional phenotype we observed upon myosin 18Aα knockdown/knockout was a significant increase in the number of primary dendrites. This phenotype could be quite interesting to pursue for two related reasons. The first has to do with the growing evidence that acentrosomal Golgi elements like Golgi outposts and Golgi satellites, which reside in primary dendrites and at dendritic branch points (outposts), as well as throughout dendritic arbors (satellites), play key roles in the morphogenesis of neuronal dendrites (Jan and Jan, 2010; Lewis and Polleux, 2012; Valenzuela and Perez, 2015; Hanus and Ehlers, 2016; Mikhaylova *et al*., 2016; Kennedy and Hanus, 2019). In addition to possessing a variety of Golgi processing enzymes, these dendritic Golgi elements commonly serve as sites of microtubule nucleation, allowing the creation of acentrosomal microtubule arrays that support their role in the local processing of ER-derived secretory cargo like transmembrane proteins (Ori-McKenney, Jan and Jan, 2012). Consistently, perturbations of either the localization or microtubule nucleating activity of Golgi outposts result in alterations in dendrite morphology. These alternations range from reduced dendritic arborization in Drosophila DA neurons (Ori-McKenney, Jan and Jan, 2012) to excessive numbers of primary dendrites in zebrafish Purkinje neurons (Tanabe *et al*., 2010) (matching the phenotype we observed here in myosin 18Aα knockdown Purkinje neurons).

The second and related reason to pursue the defect in dendrite morphology exhibited by myosin 18Aα knockdown/knockout Purkinje neurons is the published evidence that myosin 18A interacts with the resident Golgi protein GOLPH3 to promote the ribbon-like morphology of the Golgi apparatus in tissue culture cells (Dippold *et al*., 2009). Adding to the intrigue is the recent report that endogenous myosin 18A is present on Golgi outposts in oligodendrocytes (Fu *et al*., 2019). Together, these observations suggest that the defect in primary dendrite number exhibited by Purkinje neurons lacking myosin 18Aα, as well as other possible defects in dendrite morphology that might be revealed upon more extensive analyses, could be due to a defect in the localization and/or function of Golgi outposts and satellites. Interestingly, myosin 18Aα lacking its non-helical tailpiece was incapable of rescuing the defect in primary dendrite number (Figure S7), arguing that β-Pix may play a critical role in the underlying process of primary dendrite specification. Consistently, multiple studies have demonstrated that Cdc42 (one target of β-Pix’s GEF activity) plays a central role in Golgi outpost formation and trafficking (Ramakers *et al*., 2011; Friesland *et al*., 2013; Meseke, Rosenberger and Förster, 2013; Long and Simpson, 2017). Future studies should seek to elaborate upon these apparent connections between myosin 18Aα, β-Pix and dendritic Golgi elements in regulating dendrite morphogenesis.

---

**Figure S1. Myosin 18Aα is the predominant isoform expressed in the cerebellum.** Representative Western blot performed on a human Jurkat T cell extract (lane 1), a whole adult mouse cerebellum extract (lane 2), and whole adult mouse cardiac muscle extract (lane 3) probed with an anti-myosin 18A antibody against the C-terminal 300 residues of myosin 18Aα/β. The expected positions for the heavy chains of myosin 18Aα, myosin 18β and myosin 18Aγ are indicated. The origin of the large, diffuse signal in the cardiac sample, and the identity of the lower molecular weight band in that sample, are unknown.

**Figure S2. Endogenous myosin 18Aα localizes to Purkinje neuron spines in frozen tissue sections.** (A1-A6) Panels A1-A3 show a representative example of a portion of a frozen sagittal section from a perfusion-fixed adult mouse that was stained for Calbindin (A1) and myosin 18Aα (A2); Overlay (A3). Panels A4-A6 show enlargements of the boxed region in Panel A3. Arrowheads in Panel A6 point to some of the many myosin 18Aα-positive spines. The significance of the colocalization between Calbindin D28K and myosin 18Aα was confirmed by determining the Pearson’s Correlation co-efficient (see Methods). Scale bars = 500 μm (A), 50 μm (B) and 5 μm (C).

**Figure S3. Myosin 18Aα’s N-terminal F-actin binding site, but not is N-terminal PDZ domain or KE-rich region, contributes to spine targeting.** Shown are portions of dendrites from representative cultured Purkinje neurons (DIV 18) that were expressing mCherry as a volume marker (A1-D1) and one of the following mGFP-tagged proteins: myosin 18Aα-NT (WT N-terminus) (A2), myosin 18Aα-NT-ΔKE (KE-rich region deleted) (B2), myosin 18Aα NT-PDZ^mut^ (function blocking mutation in the PDZ domain) (C2), and myosin 18Aα AB^mut^ (actin binding site mutated) (D2). Panels A3-D3 show the overlaid images. (E) Shown are the fold-enrichments in the spine over the adjacent dendrite for each GFP construct, as determined by ratio imaging. The N values, which correspond to the total number of spines scored using at least 6 neurons across three independent experiments, are indicated below each measurement. Scale bar: 5 μm (D3). *** *p*<0.001; **** *p*<0.0001; n.s., not significant.

**Figure S4. Myosin 18Aα’s N-terminal extension binds F-actin.** (A) SDS-PAGE of an F-actin co-sedimentation assay using a GST-tagged version of myosin 18Aα’s N-terminal extension (NT) and the indicated ratios of NT to F-actin (obtained by fixing the concentration of NT at 1 µM and increasing the molar ratio of F-actin as indicated). S = supernatant, P = pellet. (B) As in (A) except using a GST-tagged version of myosin 18Aα’s N-terminal extension harboring a function blocking mutation in the actin binding site (residues V117 and L118 both changed to A) (NT-AB^mut^). (C) Binding isotherms for NT (closed circles) and NT-AB^mut^ (open circles) (representative of two independent experiments). The K_d_ values shown were determined by fitting the data to a one-site binding hyperbola.

**Figure S5. miRNA-mediated knockdown of myosin 18Aα.** (A) Representative Western blot of whole cell extracts of mouse NIH 3T3 mouse fibroblasts that were either untransfected (lane 1) or transfected with two different miRNAs directed against myosin 18Aα (miRNA #1, lane 2; miRNA #2, lane 3). Rack1 was used as a loading control. Densitometry indicated that the level of myosin 18Aα was reduced by 81 ± 10% and 84 ± 9% by miRNAs #1 and #2, respectively (N=3). Because 3T3 fibroblasts (unlike Purkinje neurons) also express myosin 18Aβ (which shares with myosin 18Aα both miRNA targeting sequences), its levels were also reduced by both miRNAs by ∼80%. (B1-B3) Shown is a portion of a dendrite from a representative cultured Purkinje neuron (DIV 18) that was expressing a scrambled control miRNA and that was fixed and stained for mCherry (to amplify the signal from the miRNA-associated mCherry volume marker) (B1) and for myosin 18Aα (B2); Overlay (B3). (C1-C3) As in Panels B1-B3, except the Purkinje neuron was expressing myosin 18Aα miRNA #2. Control and knockdown neurons were imaged on the same day with identical settings on a Zeiss 780 confocal microscope so that the intensities for myosin 18Aα can be directly compared. (D) Representative Western blot of whole cell extracts of mouse embryo fibroblasts (MEFs) isolated from a conditional myosin 18A knockout mouse (M18 cKO; see Methods) that were (lane 2) or were not (lane 1) transfected with a puromycin-resistant plasmid harboring Cre recombinase (Puro-Cre). The expected positions for the heavy chains of myosin 18Aα and myosin 18 Aβ are shown. Clathrin heavy chain (HC) was used as a loading control. Scale bar: 5 μm (B3).

**Figure S6. Knockout of myosin 18Aα also results in a defect in spine maturation.** (A) A representative, Calbindin-stained Purkinje neuron (DIV 18) isolated from the myosin 18A cKO mouse. (B) As in (A) except following treatment with recombinant, cell-permeable Tat-Cre on DIV 5. (C) As is (B) except the Purkinje neuron was isolated from a wild type (WT) mouse. The insets for Panels A-C show spine morphology. (D) Measurements of spine length (in µm) on DIV 18 Purkinje neurons treated as indicated (green, cKO Purkinje neuron; red, cKO Purkinje neuron plus Tat-Cre; blue, WT Purkinje neuron plus Tat-Cre). The N values, which represent scoring 6 neurons per condition from one experiment, are indicated below each measurement. (E) Measurements of spine density (in numbers per µm^2^) on DIV 18 Purkinje neurons treated as indicated. The N values, which represent scoring 6 neurons per condition from one experiment, are indicated below each measurement. Scale bar: 20 μm (A-C) and 5 μm (A-C insets). ** *p*<0.01; **** *p*<0.0001; n.s, not significant.

**Figure S7. Loss of myosin 18Aα results in an increase in the number of primary dendrites.** The number of primary dendrites per neuron (DIV 18) for the indicated samples. The number of neurons scored is indicated within each bar. * *p*<0.05; **** *p*<0.0001; n.s, not significant. We note that while mature Purkinje neurons in situ possess only one primary dendrite, our control cells typically exhibit two, as seen previously for cultured Purkinje neurons (Hirai and Launey, 2000; Tanaka et al., 2006).

## ACKNOWLEDGMENTS

This work was supported in part by the Intramural Research Program of the National Heart, Lung, and Blood Institute (NHLBI) (ZIAHL006123 to JAH) and in part by the Intramural Research Program of the National Institute on Deafness and Other Communication Disorders (NIDCD) (DC000039 to TBF; Advanced Imaging Core DC000081 to RSP). The authors thank Dr. Xufeng Wu (NHLBI Light Microscopy Core) for imaging support.

