## Supplementary figures and images for "Myosin 18Aα targets guanine nucleotide exchange factor β-Pix to dendritic spines of cerebellar Purkinje neurons to promote spine maturation"

### Supplemental Figures

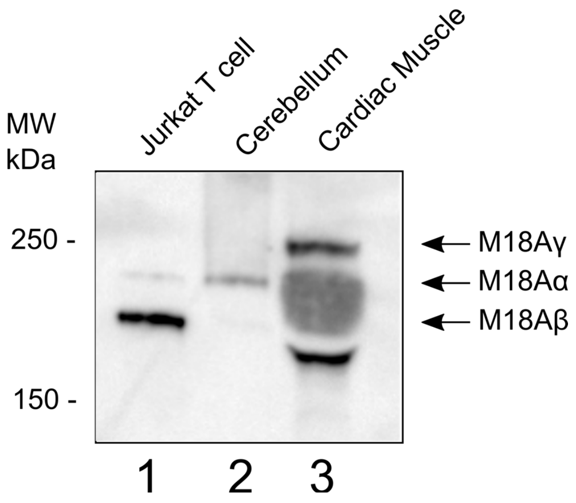

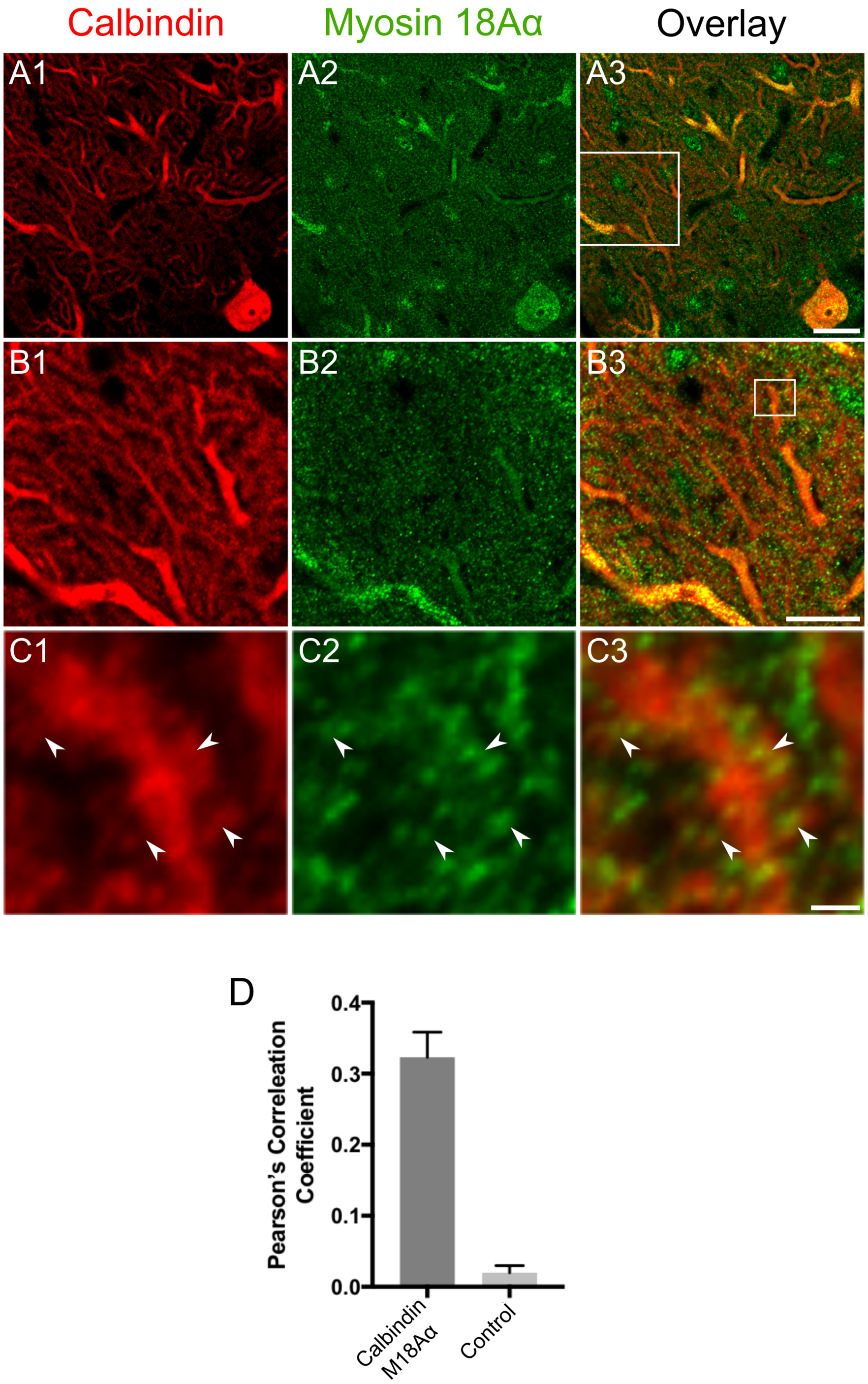

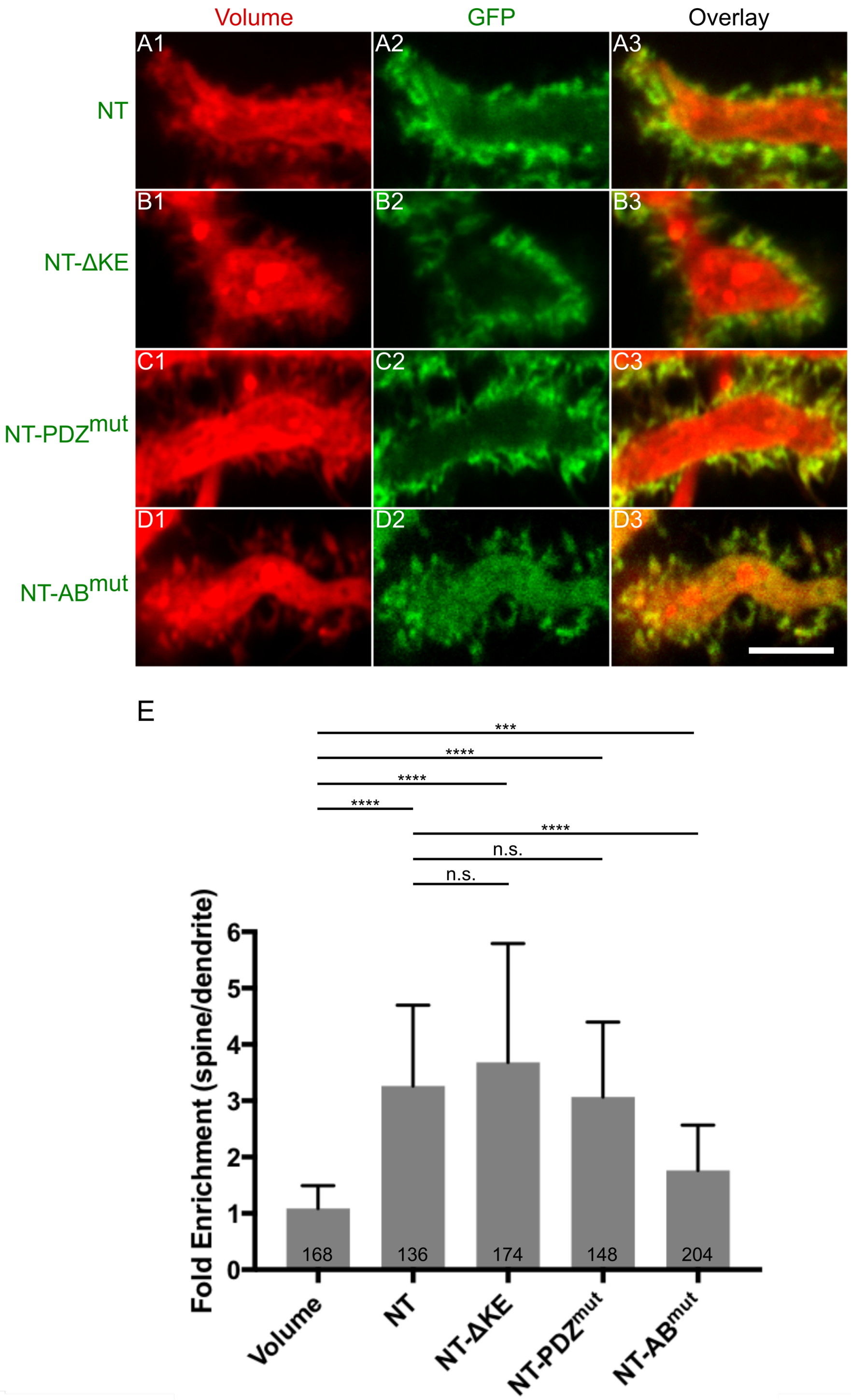

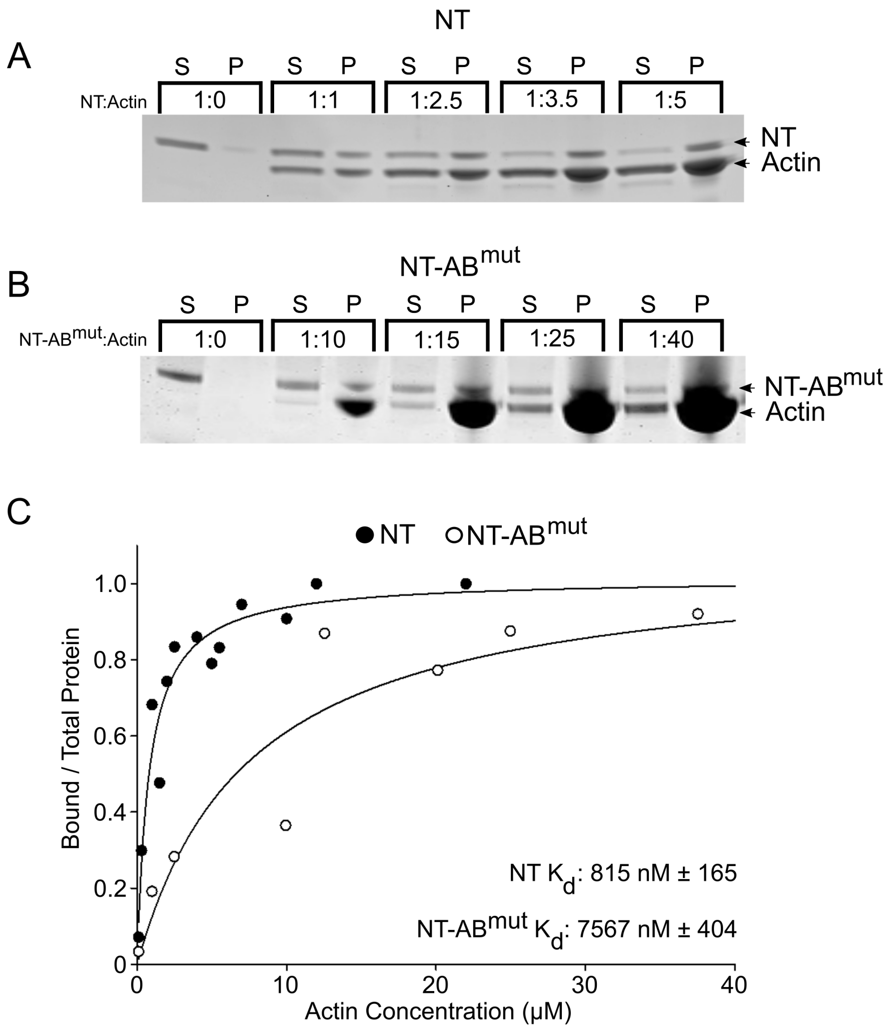

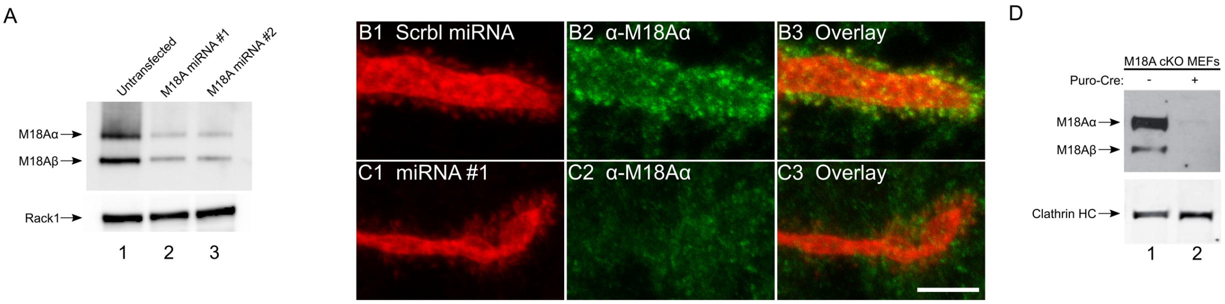

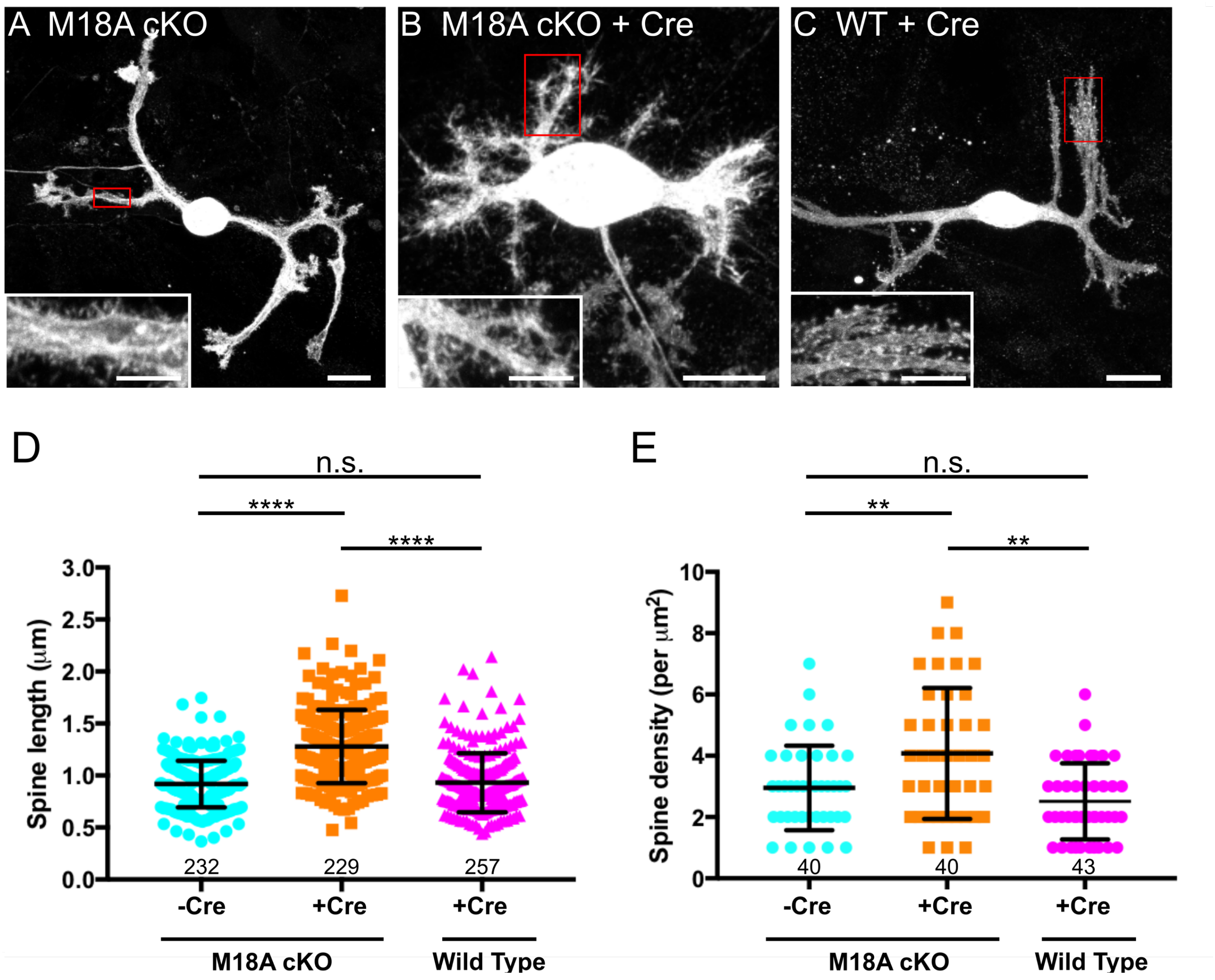

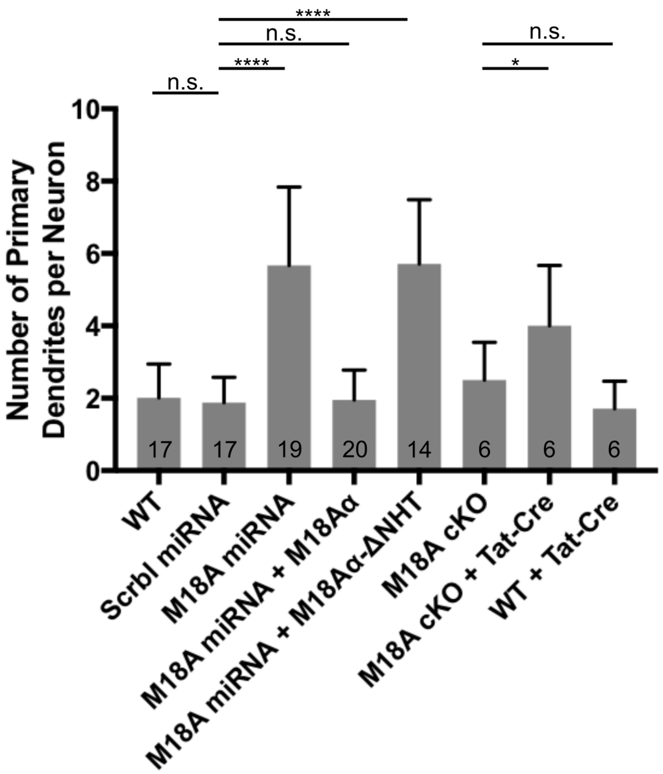
